# A post-assembly conformational change makes the SARS-CoV-2 polymerase elongation-competent

**DOI:** 10.1101/2025.01.10.632299

**Authors:** Misha Klein, Arnab Das, Subhas C. Bera, Thomas K. Anderson, Dana Kocincova, Hery W. Lee, Bing Wang, Flavia S. Papini, John C. Marecki, Jamie J. Arnold, Craig E. Cameron, Kevin D. Raney, Irina Artsimovitch, Mathias Götte, Robert N. Kirchdoerfer, Martin Depken, David Dulin

## Abstract

Coronaviruses (CoV) encode sixteen non-structural proteins (nsps), most of which form the replication-transcription complex (RTC). The RTC contains a core composed of one nsp12 RNA-dependent RNA polymerase (RdRp), two nsp8s and one nsp7. The core RTC recruits other nsps to synthesize all viral RNAs within the infected cell. While essential for viral replication, the mechanism by which the core RTC assembles into a processive polymerase remains poorly understood. We show that the core RTC preferentially assembles by first having nsp12-polymerase bind to the RNA template, followed by the subsequent association of nsp7 and nsp8. Once assembled on the RNA template, the core RTC requires hundreds of seconds to undergo a conformational change that enables processive elongation. In the absence of RNA, the (apo-)RTC requires several hours to adopt its elongation-competent conformation. We propose that this obligatory activation step facilitates the recruitment of additional nsp’s essential for efficient viral RNA synthesis and may represent a promising target for therapeutic interventions.

## Introduction

The COVID-19 pandemic is caused by the severe acute respiratory syndrome coronavirus 2 (SARS-CoV-2) with devastating impacts on public health that are still ongoing (WHO: https://data.who.int/dashboards/covid19/cases?n=c). Novel CoVs may also emerge in the future, as the present pandemic comes close on the heels of the MERS-CoV in 2012 and SARS-CoV-1 in 2002. Viral genome replication and transcription are important targets for therapeutic interventions due to their conserved nature across CoVs, and treatments based on nucleotide analogs, such as remdesivir (1) and molnupiravir (2), have proven to be effective. However, CoVs have a great ability to evolve and adapt, either via mutation or recombination (3–5). Having a precise understanding of the molecular determinants in CoV replication and transcription will help design future antiviral drugs to counter such adaptation.

CoVs are positive (+)RNA viruses with a ∼30 kb long single-stranded (ss) genome. The 5’-proximal two thirds encode for the sixteen non-structural proteins (nsp(s)) (13). These nsps are encoded on two open reading frames separated by a ribosomal frameshifting sequence within the nsp12-polymerase gene. Translation produces two polyproteins: one spanning from nsp1 to nsp11, and the other from nsp1 to nsp16. These polyproteins are subsequently processed into individual nsps by two proteases—nsp3 (papain-like protease) and nsp5 (3C-like protease). Notably, nsp5 processes all nsps from nsp4 to nsp16 (14). Most of the nsps associate to form the replication-transcription complex (RTC), which synthesizes all viral RNAs in the infected cell (15, 16). The minimal complex capable of RNA synthesis consists of the nsp12-polymerase associated with nsp7 and nsp8 in a 1:1:2 stoichiometry (hereafter ‘core RTC’) (10, 17–20). Nsp7 and nsp8 are required for *in vitro* RNA primer-extension by the core RTC (21–24).

Previous studies have shown that it takes several minutes for the CoV core RTC to perform detectable primer-extension in *in vitro* bulk assays (23, 24). This is particularly striking, as the SARS-CoV-2 core RTC is the fastest RNA polymerase reported to date, with a nucleotide addition rate up to ∼180 nt/s at 37°C (11). The apparent lag time in primer-extension assays therefore seems to stem from a step prior to RNA synthesis and has been proposed to arise from a stepwise assembly of the individual nsps into the core RTC (22, 23, 25). However, no quantitative assessment of different contributors to the lag times has been reported to date. Characterizing the mechanism through which nsp7 and nsp8 regulate primer-extension by the core RTC requires a more in-depth comparison of their influence, both before (hereafter ‘activation phase’) and during (hereafter ‘elongation phase’) RNA synthesis (**Figure 1A**).

**Figure 1:**
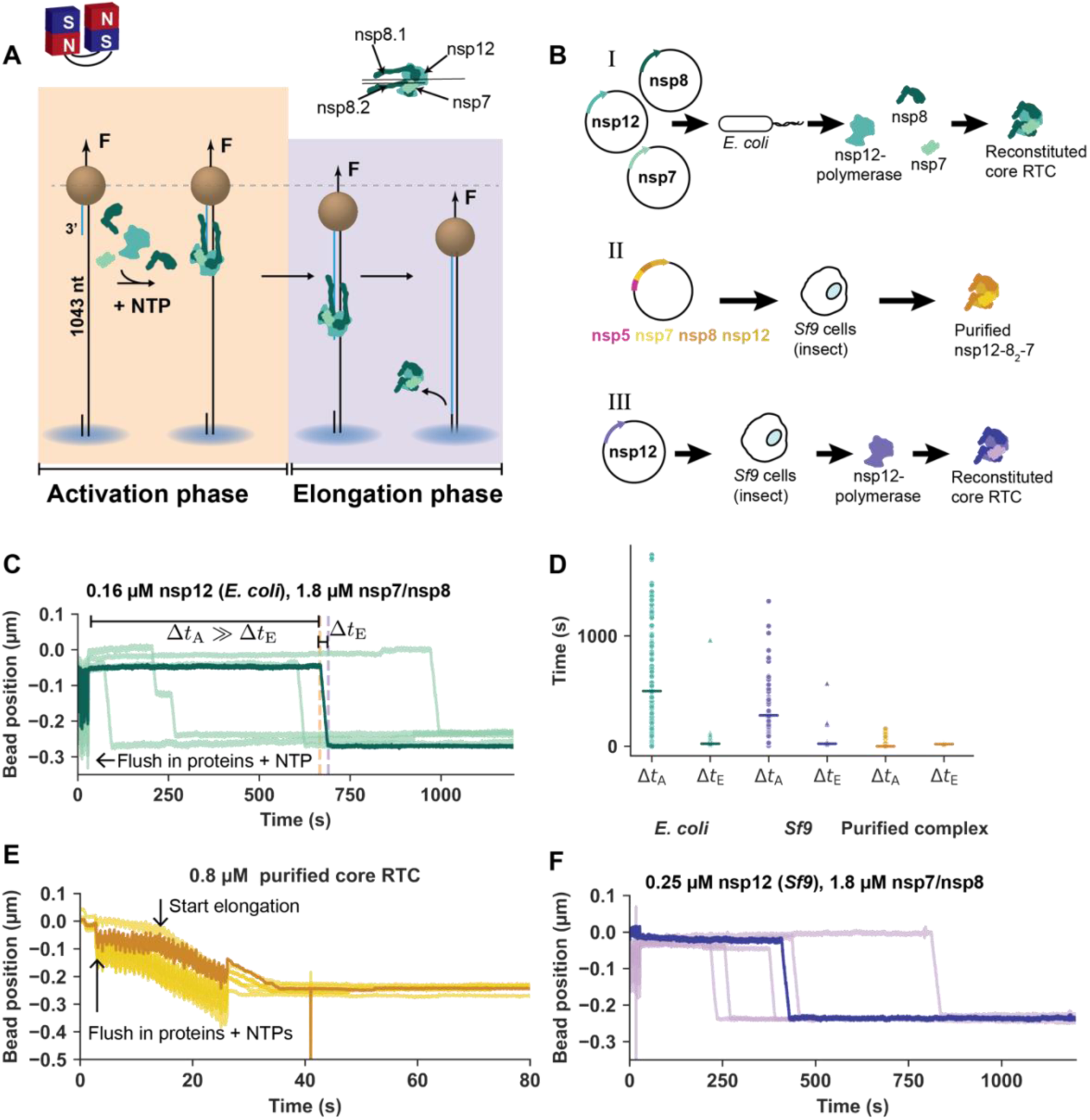
Slow activation by reconstituted SARS-CoV-2 core RTCs. **(A)** Schematic of the magnetic tweezers assay to monitor activation and elongation by the SARS-CoV-2 core RTC (**Materials and Methods**). **(B)** Schematic representation different methods of obtaining the core RTC used throughout this work. (I) nsp7, nsp8 and nsp12-polymerase were separately and recombinantly expressed in *E. coli*, (II) purification of core RTC complex after expressing a bacmid containing nsp5-Mpro (main protease), nsp8, nsp7, and nsp12-polymerase in *Sf9* cells, (III) nsp7, nsp8 and nsp12-polymerase were separately and recombinantly expressed in *E. coli and* nsp12-polymerase was recombinantly expressed in *Sf9* (**Materials and Methods**). Color coding for the different core RTCs is kept in subsequent figures to represent to related data. **(C)** Example time traces for the reconstituted SARS-CoV-2 core RTC using 0.16 µM nsp12-polymerase (see (I) in panel B) and 1.8 µM of nsp7 and nsp8. The orange and purple dashed lines indicate the end of the activation and elongation phases. Δ*t*_A_ and Δ*t*_E_ are their respective durations. **(D)** Δ*t*_A_ and Δ*t*_E_ for all recorded traces in experiments of which representative subsets are shown in panels C, E and F. Horizontal markers indicate the group medians. **(E)** Example time traces using 0.8 µM of the purified SARS-CoV-2 core RTC (see (II) in panel B). **(F)** Example time traces for the reconstituted SARS-CoV-2 core RTC using 0.25 µM nsp12-polymerase expressed in *Sf9* (see (III) in panel B) and 1.8 µM of nsp7 and nsp8.

**Figure 2:**
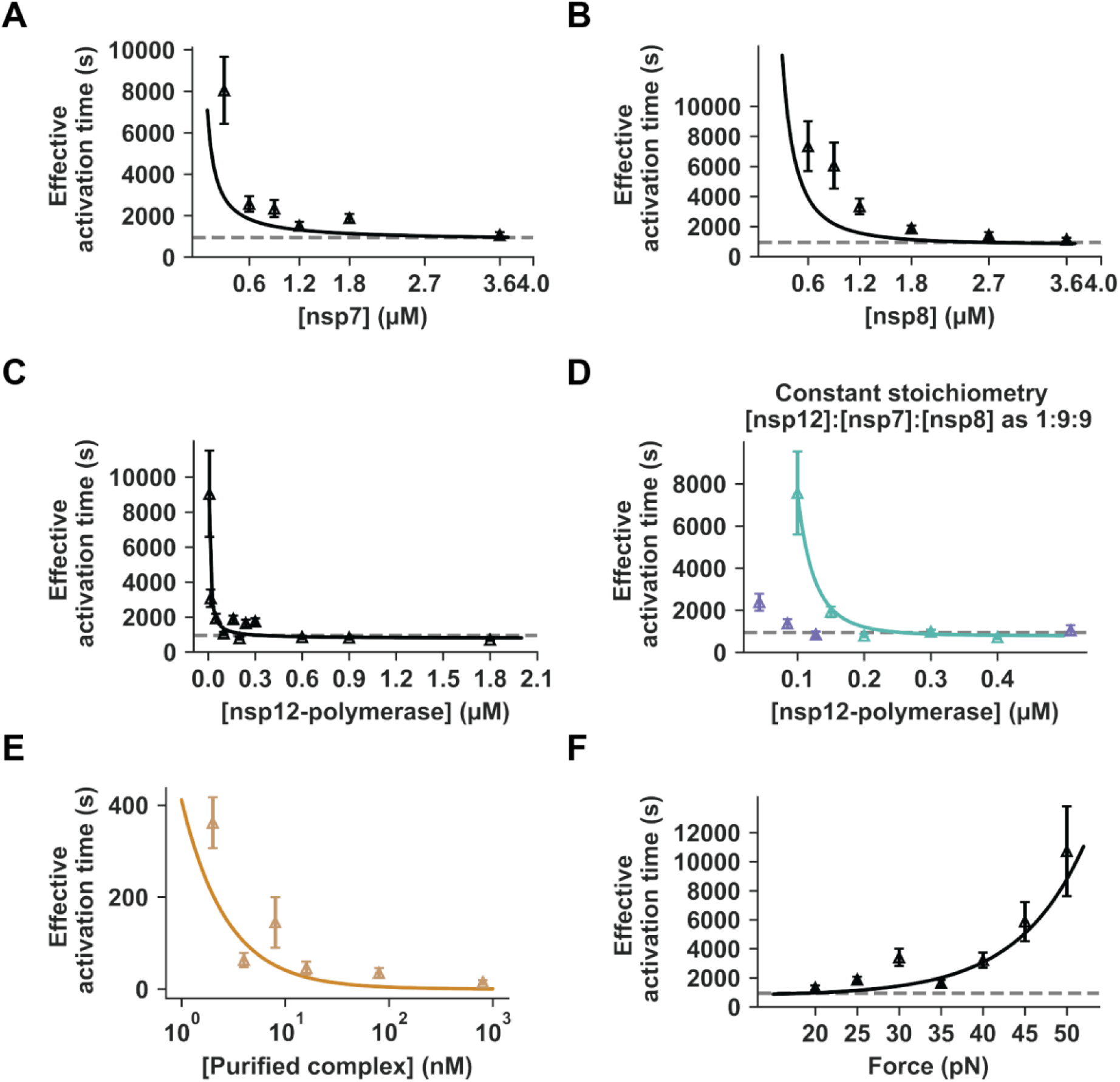
Assembly and RNA binding of the core RTC cannot fully account for the observed activation times. When not being varied, experiments were performed using 0.16 µM of nsp12-polymerase, 1.8 µM nsp7, 1.8 µM nsp8, and 25 pN. Effective activation times across **(A)** nsp7 concentration, **(B)** nsp8 concentration, **(C)** nsp12-polymerase concentration (expressed in *E. coli*). **(D)** Activation times versus nsp12-polymerase concentration, while maintaining a constant stoichiometry of [nsp12-polymerase]:[nsp7]:[nsp8] as 1:9:9 in solution. Core RTC reconstituted using nsp12-polymerase expressed in *E. coli* (turquoise) or *Sf9* (purple). **(E)** Activation times versus concentration of purified complex. **(F)** Activation times versus force (reconstituted core RTC, all proteins expressed in *E. coli*). Error bars represent 95% confidence intervals estimated through bootstrapping (**Materials and Methods)**. Effective activation times shown incorporate the fraction of events lasting longer than the recording (**Materials and Methods**). Solid curves are the best fit of our mechanochemical model (**Materials and Methods, Figure 5 and Figure S6**). Dashed horizontal lines are shown to guide-the-eye.

We set out to understand the path the CoV core RTC proteins take towards polymerase activity by using a high-throughput single-molecule magnetic tweezers assay (26) in conjunction with kinetic modelling. Previously, we used the same assay to study the elongation phase of the SARS-CoV-2 core RTC in-depth (26). Here, we used our ability to detect the onset of elongation from individual RNA primer-templates to differentiate between the time spent by single core RTCs on activation and elongation. Our data and kinetic model reveal that a core RTC is not in an elongation-competent conformer upon assembly. As such, the core RTC must undergo a conformational change that allows it to enter the elongation phase. We start by confirming that a core RTC (reconstituted from individually expressed proteins) exhibits a considerable lag-time prior to elongation (23, 24). The lag time does not stem from the time required to assemble the complex since activation times remain substantial at saturating protein concentrations. Instead, nsp7, nsp8, and the template RNA appear to greatly limit the conformational space to be explored by the core RTC, allowing it to adopt its elongation-competent conformer within minutes. The conformational change occurs prior to the elongation phase as factors influencing the kinetics of the activation phase have no effect on the elongation phase. Finally, even in the presence of nsp13-helicase an average activation time of several hundreds of seconds remains. Our results reveal a mandatory step towards assembling a complete and functional CoV-RTC, opening new avenues for antiviral drug development.

## Materials and Methods

### Purification and recombinant protein expression of nsp7 and nsp8 from SARS-CoV-2

This protocol was described in detail in (6, 26). The SARS-CoV-2 nsp7 and nsp8 genes were cloned into pET46 (Novagen) with an N-terminal 6 histidine tag and a TEV protease site for purification. The proteins were recombinantly expressed in Rosetta2 pLys *E.coli* (Novagen) and purified using Ni-NTA agarose beads. Nsp7 and nsp8 proteins were further purified by size exclusion chromatography. Purified proteins were dialyzed into storage buffer (10 mM HEPES, pH 7.4, 300 mM NaCl, 2 mM DTT), aliquoted, and stored at –80 °C.

### Purification and recombinant protein expression of nsp12-polymerase in *Sf9* cells

This protocol was described in detail in (6, 26).The SARS-CoV-2 nsp12-polymerase gene was cloned into pFastBac with a TEV site and strep tag (Genscript) for efficient purification. The bacmid was created in DH10Bac *E.coli* (Life Technologies) and amplified in *Sf9* cells (Expression Systems) with Cellfectin II (Life Technologies) to generate recombinant baculovirus. Sf21 cells were infected, and the cell lysate cleared by centrifugation and filtration. The protein was purified via strep tactin agarose and further polished in size exclusion chromatography. Purified protein was dialyzed into storage buffer (10 mM HEPES, pH 7.4, 300 mM NaCl, 0.1 mM MgCl_2_, 2 mM TCEP), aliquoted, and stored at –80 °C.

### Purification and recombinant protein expression of nsp12-polymerase in *E. coli*

The protocol used to obtain nsp12-RdRp used for these experiments was decribed in detail in (27). Nsp12-polymerase was overexpressed in *E. coli* BL21(DE3) cells (Novagen). The cleared lysate was applied to Ni21-NTA resin (Cytiva) and the eluted protein was further purified by anion exchange chromatography and size exclusion chromatography. Purified protein was dialyzed into storage buffer (20 mM HEPES, pH 7.5, 150 mM KCl, 45% glycerol, 1 mM MgCl_2_, 1 mM DTT), aliquoted, and stored at –80 °C.

### Purification of co-translated SARS-CoV-2 core RTC

The protocol was described in detail in (12). The pFastBac-1 (Invitrogen, Burlington, ON, Canada) plasmid with the codon-optimized synthetic DNA sequences (GenScript, Piscataway, NJ, USA) coding for SARS-CoV-2 (NCBI: QHD43415.1) nsp5, nsp7, nsp8, and nsp12 were used as a starting material for protein expression in insect cells (*Sf9*, Invitrogen, Waltham, MA, USA) (8, 9, 12, 28). We employed the MultiBac (Geneva Biotech, Indianapolis, IN, USA) system for protein expression in insect cells (Sf9, Invitrogen) according to published protocols (29, 30). SARS-CoV-2 protein complexes were purified using nickel-nitrilotriacetic acid affinity chromatography of the nsp8 N-terminal 8-histidine tag according to the manufacturer’s specifications (Thermo Fisher Scientific, Waltham, MA, USA). Purified protein was dialyzed into storage buffer (100 Tris-HCl (pH 8), 1000 mM NaCl, 5 mM TCEP, 0.01 % Tween-20, 200 mM imidazole, and 40% glycerol), aliquoted and stored at –20 °C.

### Purification and recombinant protein expression of nsp13-helicase

The coding sequence for nsp13 from the SARS CoV-2 Washington isolate (Genbank MN985325) was synthesized as an *E. coli* codon-optimized fragment (GenScript, Piscataway NJ) and cloned into the *Bsa*I site of the pSUMO plasmid (LifeSensors, Malvern, PA) to produce an *N*-terminal six histidine-tagged SUMO-nsp13 fusion cassette (6XHis-SUMO-nsp13). Final plasmids were sequence-verified through the UAMS Sequencing Core Facility using a 3130XL Genetic Analyzer (Applied Biosystems, Foster City, CA). The SUMO-nsp13 construct was transformed into Rosetta2 cells, and colonies were grown overnight at 37°C in NZCYM (Research Products International, Mount Prospect, IL) supplemented with kanamycin (50 µg/ml) and chloramphenicol (25 µg/ml). The cultures were diluted 1:100 into fresh antibiotic-containing NZCYM media and grown to an OD_600 nm_ of 0.8-1. The bacterial media was supplemented with 0.1 mM ZnSO_4_ and 0.2% dextrose and cooled on ice for 10 min. Protein expression was induced with 0.2 mM isopropyl β-D-1-thiogalactopyranoside (IPTG) at 18°C for 12-16 hours. The cells were harvested by centrifugation at 4,000 x *g* for 15 min at 4°C, and pellets stored at –80°C.All purifications steps were carried out on ice or at 4°C. Pellets were resuspended in lysis buffer (50 mM sodium phosphate, pH 8.0, 300 mM NaCl, 1 mM β-mercaptoethanol, 10% glycerol and 20 mM imidazole) supplemented with 2 mM phenylmethylsulfonyl fluoride (PMSF) and 1X EDTA-free protease inhibitor cocktail (Pierce). Bacteria were lysed by microfluidization and the lysate clarified by centrifugation at 17,000 x *g* for 1 hour at 4°C. The His-tagged SUMO-nsp13 was passed through a HisTrap FF column (Cytiva) equilibrated in lysis buffer at 1 ml/minute using a Cytiva Akta FPLC. The affinity resin was washed with 20 column volumes of lysis buffer, and the protein eluted with 10 column volumes of lysis buffer containing 200 mM imidazole. The pooled SUMO-nsp13-containing fractions were dialyzed overnight into two changes of 20 mM imidazole-containing lysis buffer, and the SUMO tag cleaved with ULP-1 for 4 hours at 4°C. Digestion was confirmed by SDS-PAGE analysis. The His6-ULP-1 and His6-SUMO proteins were separated from the native nsp13 with a second round of Ni^2+^-affinity chromatography as before. Nsp13-containing flow-thru fractions were pooled, dialyzed overnight against two changes of low salt buffer (50 mM sodium phosphate, pH 6.8, 150 mM NaCl, 4 mM β-mercaptoethanol, 0.5 mM EDTA and 10% glycerol) and passed through a HighTrap SP (Cytiva) cation exchange column. Under these conditions, nsp13 did not adhere to the SP column and the flow-thru was collected in multiple fractions. The nsp13 was concentrated with an Amicon Ultra-15 centrifugation filter unit to a volume of ∼1.5 mls and loaded on to a Sephacyl S200-HR HiPrep 26/60 column (Cytiva) equilibrated with nsp13 Storage Buffer (25 mM HEPES, pH 7.5, 150 mM NaCl, 0.5 mM TCEP and 20% glycerol). The final nsp13 was quantified by UV spectrophotometry at 280 nm using the expected extinction coefficient of 68,785 M^-1^ cm^-1^ and confirmed using the BCA Protein Assay (Pierce). Protein samples were aliquoted, flash frozen, and stored at –80°C.

### RNA hairpin fabrication

The fabrication of the RNA hairpin has been described in detail in (31). The RNA hairpin is made of a 499 bp double stranded RNA stem terminated by a 20 nt loop that is assembled from three ssRNA annealed together (**Figure S1A**). The RNA stem is flanked by two spacers, ∼800 bp each, containing a biotin– and a digoxigenin-handle, respectively. A gap of 25 nt between the biotin-handle and the hairpin stem serves as the loading site for the polymerase (**Figure S1A**). At forces above 22 pN, the hairpin opens and frees up a 1043 nt ssRNA template for the SARS-CoV-2 polymerase (**Figure S1B**). The RNA construct was synthesized by amplifying DNA fragments in PCR and *in vitro* transcribing them (NEB HiScribe T7 High Yield RNA Synthesis Kit) after purification (Monarch PCR and DNA cleanup kit). ssRNA fragments containing biotin-or digoxigenin-labels were synthesized with biotin-UTP or digoxigenin-UTP (Jena Biosciences) in the reaction. Transcripts were mono-phosphorylated (Antarctic Phosphatase and T4 Polynucleotide Kinase), annealed and ligated. The RNA template sequence is provided in (26).

### Flow cell assembly and surface functionalization

The fabrication procedure for flow cells has been described in detail in (32). To summarize, we sandwiched a double layer of Parafilm by two #1 coverslips, the top one having one hole at each end serving as inlet and outlet, the bottom one being coated with a 0.1% m/V nitrocellulose dissolved in amyl acetate solution. The flow cell is mounted into a custom-built holder and rinsed with ∼1 ml of 1x phosphate buffered saline (PBS) solution. 3 µm diameter polystyrene reference beads are attached to the bottom coverslip surface by incubating 100 µL of a 1:1000 dilution in PBS (LB30, Sigma Aldrich, stock conc.: 1.828*1011 particles per milliliter) for ∼3 min. The tethering of the magnetic beads by the RNA hairpin construct relies on a digoxygenin/anti-digoxygenin and biotin-streptavidin attachment at the coverslip surface and the magnetic bead, respectively. Therefore, following a thorough rinsing of the flow cell with PBS, 50 µL of anti-digoxigenin (50 mg/mL in PBS) is incubated for 30 min. The flow cell was rinsed with 1mL of 10 mM Tris, 1 mM EDTA pH 8.0, 750 mM NaCl, 2 mM sodium azide buffer to remove excess of anti-digoxigenin followed by rinsing with another 0.5 ml of 1x TE buffer (10 mM Tris, 1 mM EDTA pH 8.0 supplemented with 150 mM NaCl, 2 mM sodium azide). The surface is then passivated by incubating bovine serum albumin (BSA, New England Biolabs, 10 mg/ml in PBS and 50% glycerol) for 30 min and rinsed with 1x TE buffer.

### Single-molecule SARS-CoV-2 primer-extension experiments

20 µL of streptavidin coated Dynabeads M-270 magnetic beads (ThermoFisher) were mixed with ∼0.1 ng of RNA hairpin (total volume 40 µL) and incubated for ∼5 min before rinsing with ∼2 ml of 1x TE buffer to remove any unbound RNA and the excess of magnetic beads. RNA tethers were sorted for functional hairpins by looking for the characteristic jump in extension of the correct length (∼ 0.6 µm at 22 pN) due to the sudden opening of the hairpin during a force ramp experiment (**Figure S1C)** (26, 31). The flow cell was subsequently rinsed with 0.5 ml reaction buffer (50 mM HEPES pH 7.9, 10 mM DTT, 2 mM EDTA, 5 mM MgCl2). After starting the data acquisition, the hairpin tether quality was tested by ramping the force up to monitor the typical cooperative opening signature of a hairpin, i.e. a vertical jump of the magnetic bead by ∼0.6 µm when reaching a critical opening force of ∼22 pN (**Figure S1D**). The force was subsequently decreased to and maintained at 25 pN, unless specified otherwise. 100 µL of reaction buffer containing the proteins and 500 µM of {A,U,C,G}TP (Jena Biosciences) were flushed into the flow chamber (**Figure 1A**). SARS-CoV-2 core RTC activity traces were spotted as a downward movement of the bead, indicating the conversion of the ssRNA template into dsRNA, and therefore a shortening of the tether (**Figure 1A**). The recordings lasted 30 min. A temperature of 25 °C was maintained during all experiments. A custom written LabView routine controlled the data acquisition and the (x-, y-, z-) positions analysis/tracking of both the magnetic and reference beads (Sigma) in real-time (33). Mechanical drift correction was performed by subtracting the position of the reference bead from that of the magnetic beads, and further stabilized by an automated focusing routine that adaptively moves the objective to keep the reference bead at the same focal plane (26). The camera frame rate was fixed at 58 Hz.

### Single-molecule experiments after pre-incubating proteins

For the experiments performed in **Figure 3A and S3**, 20 µM of nsp12 (of either expression system) was added to 180 µM of both nsp8 and nsp7. Next, the mixture was placed on a heating block at 25 °C for 4.5 hrs (nsp12-polymerase expressed in *E. coli*) or 6.5 hrs (nsp12-polymerase expressed in *Sf9*). After the specified waiting time, the proteins were diluted in reaction buffer to a final concentration of 0.2 µM of nsp12-polymerase and 1.8 µM of both nsp8 and nsp7. Finally, we proceeded by performing the primer-extension experiment as described above.

**Figure 3:**
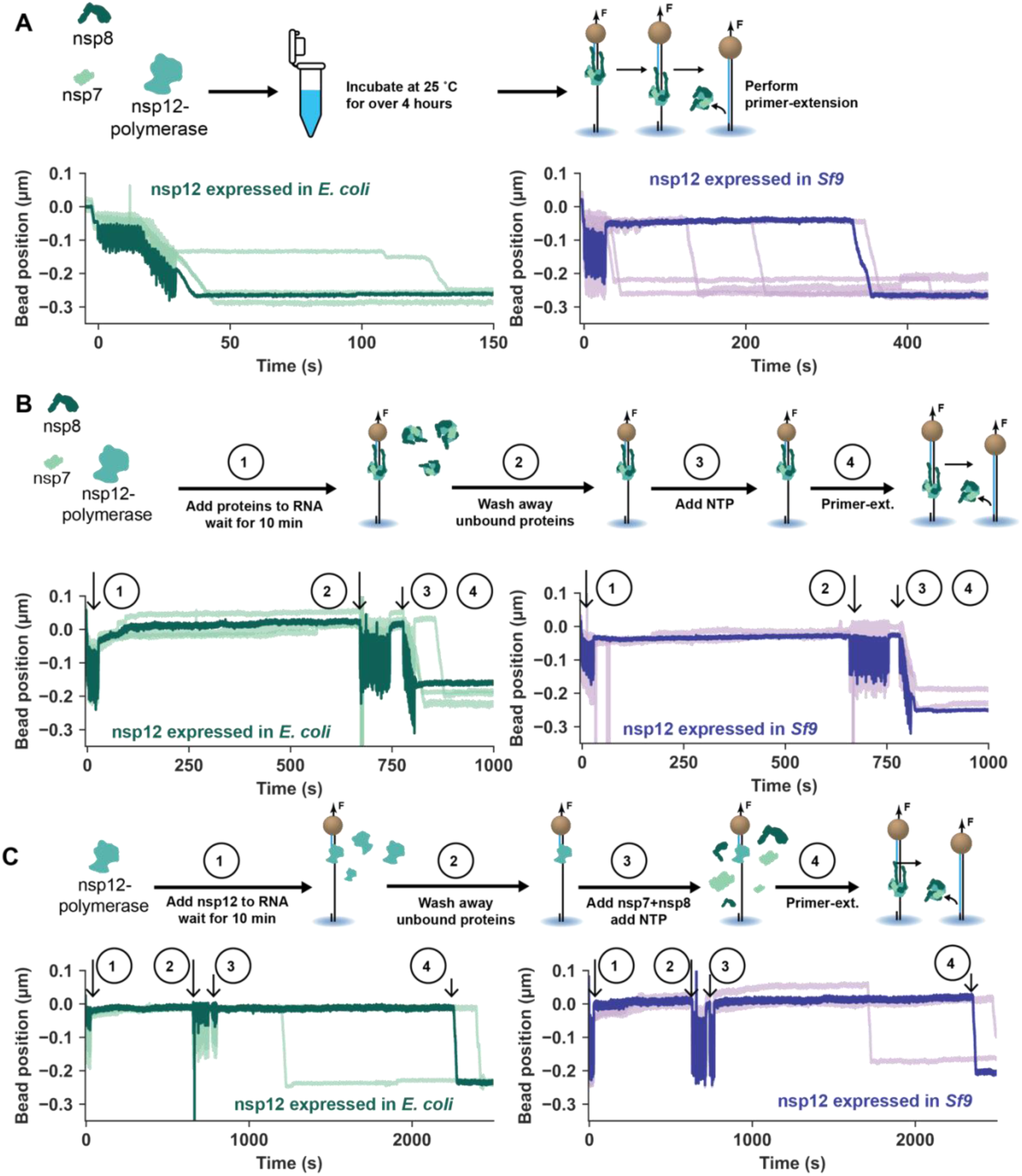
Presence of the template RNA reduces activation times. Example time traces are shown for a reconstituted core RTC. **(A)** Nsps were added together 4.5 hrs (nsp12-polymerase expressed in *E. coli*) or 6.5 hrs (nsp12-polymerase expressed in *Sf9*) prior to diluting them to 0.2 µM nsp12, 1.8 µM nsp7 and nsp8 and performing primer-extensions. **(B)** (1) The proteins were added to the RNA tethering the magnetic beads in the flow chamber. (2) Ten minutes later 500 µL of reaction buffer was used to rinse the flow chamber from free-floating proteins. (3) Finally, ribonucleotides were added and (4) Activation typically started during step (3). **(C)** (1) After tethering the magnetic beads to the RNA, 0.2 µM of nsp12-polymerase and 500 µM of NTP were added. (2) Ten minutes later 500 µL of reaction buffer was used to remove free-floating proteins. (3) The nucleotides were reintroduced together with 1.8 µM of nsp7 and nsp8. (4) RTC elongation activity.

### Single-molecule experiments after pre-incubating proteins with RNA

After forming the RNA tethers as described above, proteins were added at specified concentration to the flow chamber. The recording was started. After 10 min we removed any access of proteins in solution by rinsing the flow chamber with 500 µL of reaction buffer. Finally, primer-extension experiments were performed after the system was complemented with the missing reagents (**Figures 3BC, S4-5**). In **Figures 3B, S4**, and **S5** the proteins were incubated together with 500 µM of NTP to further confirm no activity can occur unless all nsps are present.

### Data processing of RNA synthesis traces (elongation phase)

The change in extension in micron, resulting from the ssRNA to dsRNA conversion, was subsequently converted into replicated nucleotides, low-pass filtered at 2 Hz and the dwell times were extracted using a window of 10 nt as described in (11, 26, 34, 35). Elongation times and product lengths were inferred from the final datapoint before the trajectory ended of the filtered traces (**Figure 4A**). Product lengths shorter than the full length of the template were only considered if the bead stopped moving downwards before the end of the recording. If the tether broke before the full product was synthesized (seen as the loss of the bead), that event was excluded from both elongation time and product length measurements.

**Figure 4:**
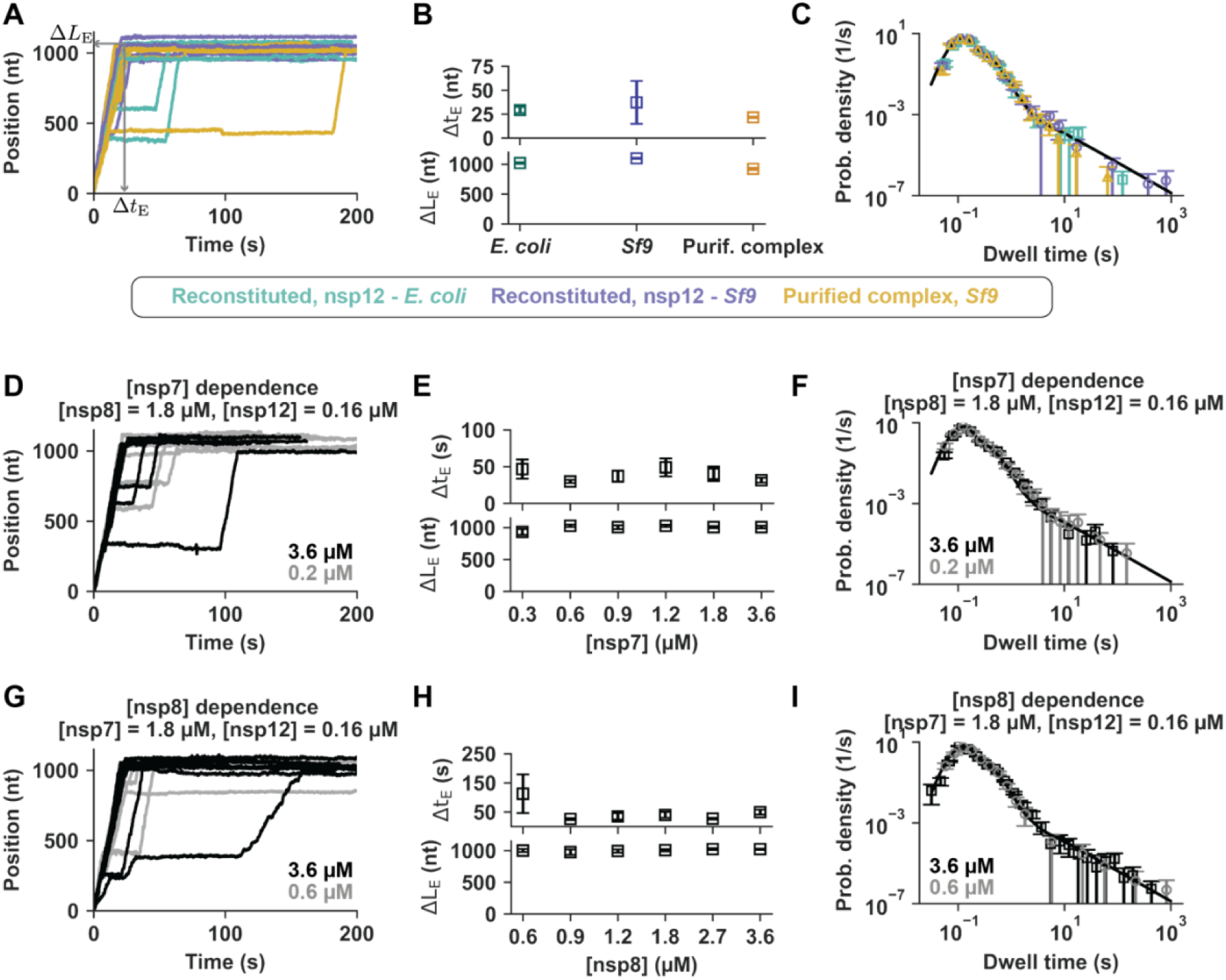
The activation phase and the mode of core RTC proteins expression do not influence the kinetics of the elongating core RTC. **(A)** Expample time traces during elongation phase of either reconstituted core RTC with 0.16 µM nsp12-polymerase expressed in *E. coli*, and 1.8 µM nsp7 and nsp8 (turquoise), reconstituted core RTC with 0.2 µM nsp12-polymerase expressed in *Sf9* and 1.8 µM nsp7 and nsp8 (purple), or 2 nM of co-translated core RTC (yellow). **(B)** Mean elongation times (Δ*t*_E_) and product lengths (Δ*L*_E_), and **(C)** dwell time distributions for the entire set of activity traces recorded. The solid line in panel C represents the distribution we previously reported in Ref. (26). **(D)** Example time traces during elongation phase of a reconstituted core RTC (nsp12-polymerase expressed in *E. coli*) at 1.8 µM (black) and 0.2 µM (grey) of nsp7. **(E)** Mean elongation times and product lengths, and dwell time distributions across nsp7 concentrations. **(G)** Example time traces during elongation of reconstituted core RTC (nsp12-polymerase expressed in *E. coli*) at 1.8 µM (black) and 0.6 µM (grey) of nsp8. **(H)** Mean elongation times and product lengths, and **(I)** dwell time distribution across nsp8 concentrations. All error bars represent 95% confidence intervals determined as described in **Materials and Methods**.

### Data processing of activation times

When RNA extension could be observed in the trace, the activation time was taken as the time between ending the addition of reagents into the flow chamber and the first datapoint when the bead starts moving downwards (**Figure 1C**). Only tethers with an open hairpin before addition of the reagents were considered (**Figure S1D**). Polymerase activity when the elongation phase started while still adding reagents was clearly recognized (**Figure 1E**), but the start time/position of these events could not be determined with high accuracy and these were assigned an activation time of zero seconds. Tethers that remained until the end of the recording were considered to determine the fraction of tethers with activity (**Figures S2, S7, and S8**). This showed that a considerable portion of traces showed no sign of RNA extension during the 30 min recordings. Assuming that activation on different tethers are independent, the activation efficiency results from a Bernoulli trial. The experimental estimate (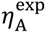) of the primer-extension efficiency (η_A_) is the total number of tethers on which we record activity (N_rec_) over to total number of quality RNA hairpin tethers (N_HP_)

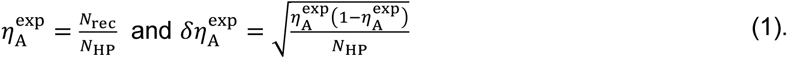

We used that 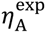 is an unbiased estimator of η_A_, and plot the 95% confidence intervals (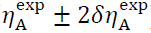) in **Figure S2**.

To extract the maximum likelihood estimator of the single-exponential time constant that best describes the distribution of activation times we included events missed due to the limited experimental time. This method has been described in detail in (36). Briefly, the maximum-likelihood estimator (MLE) for the time constant in an single-exponential distribution would equal the average of all recorded times (**Figure S2**),

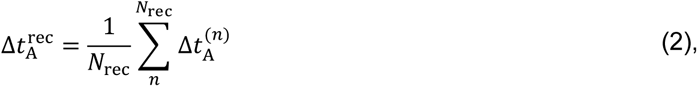

when there is no limit on the experimental observation time. In the above, 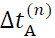 corresponds to the activation time recorded on the *n*^th^ out of the N_rec_ traces that initiate. With a finite experimental time N_cut_ traces will not initiate within the experimental time *T*_cut_, and the MLE that accounts for this is given by

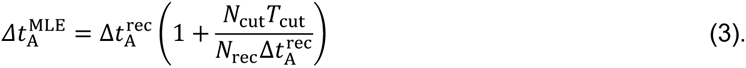

To obtain an error estimate for 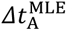, 1000 bootstrap samples were drawn from the collection of N_HP_ = N_rec_ + N_cut_ traces. In this manner we get a distribution of estimated activation times (**Equation 3**). The resulting 95% confidence intervals were used as the error estimates shown in **Figure 2**. In **Figures S7 and S8** we overlay the distributions of recorded activation times with the exponential distribution with time constant **Equation 3** and total probability 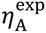.

The MLE of **Equation 3** was only applied to the experiments involving the individually purified nsps. At saturating concentrations of the purified complex, activation typically started before we stopped flushing reagents into the flow chamber (**Figure 1E** and **Figure 2E**). Yet, not all RNA tethers got converted into dsRNA during our recording (**Equation 1, Figure S2J**). The measured activation efficiencies (**Equation 1, Figure S2J**) showed little to no dependence on the concentration of the complex in this case, indicating that they do not originate in events missed due to our limited observational time; consequently we exclude these from our considerations. **Figure 2E** and **Figure S2I** therefore show the MLE of **Equation 2**. Error estimates are 95% confidence intervals from 1000 bootstrap samples drawn from the N_rec_ recorded times.

### Modeling the activation time of the reconstituted core RTC

The reaction schema underlying our kinetic model is shown in **Figure S6**. We split the activation time into two parts; 1) a ‘slow’ process that includes the nsp7– and nsp8-dependent ‘activation step’ (**Figures 5 and S6**), and 2) the time spent by the core RTC after being stabilized into an elongation-competent conformation, but before incorporation of the first nucleotide (**Figure S6**)

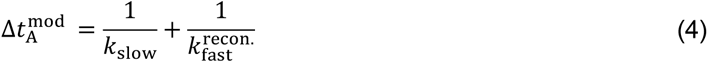

**Figure 5:**
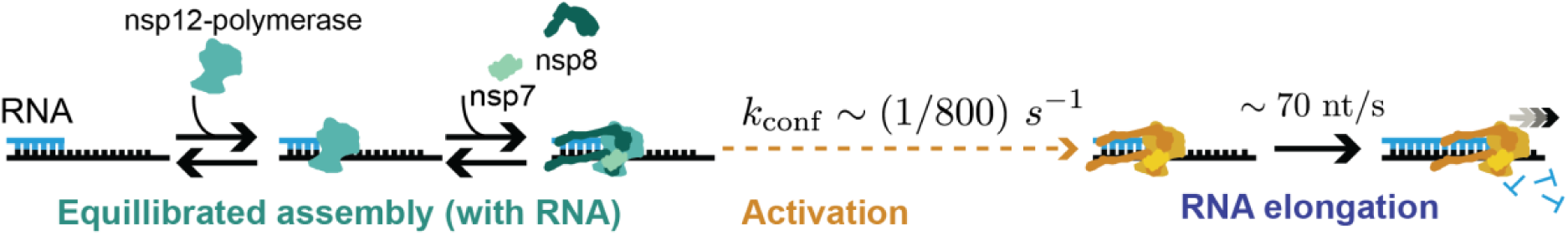
The activation of the core RTC into a processive polymerase is rate-limited by a slow conformational change following assembly. Nsp12-polymerase rapidly binds to the RNA and thereafter recruits the co-factors nsp7 and nsp8, followed by a slow and irreversible conformational change that activates the core RTC into processive elongation.

We will start by describing the rate-limiting transition (*k*_slow_). Our results showed activation is orders of magnitudes faster in the presence of the RNA template (**Figures 3AB and S3**), hence we limit the activating conformational change to the RNA bound state. With this simplification the rate of the ‘slow’ step in **Equation 4** equals the equillibrated fraction *P*_bnd_of tethers with an assembled core RTC, multiplied by the intrinsic rate of undertaking the conformational change (*k*_conf_)

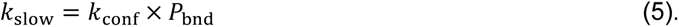

Let *c*_7_, *c*_8_ and *c*_12_ denote the concentrations of nsp7, nsp8 and nsp12-polymerase respectively. Given core RTC assembly and RNA binding happen before the rate limiting step, their respective probabilities of occurance should be equilibrated, i.e. satisfy the law of mass action and detailed balance,

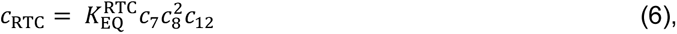

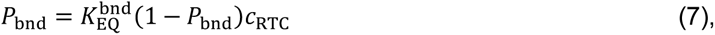

where *P*_bnd_ is the fraction of the RNA tethers with an assembled core RTCs bound to it. We also introduced the equilibrium constants for assembling the core RTC 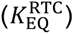 and binding the core RTC to the RNA 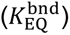. The latter having a force-dependence as shown below. Given the large disparity in apparent saturating concentrations using purified versus reconstituted core RTCs (**Figure 2**), we assumed the total number of free nsps that complex remains small enough to not effect their concentration in bulk, i.e. 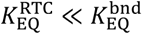. While losing some generality, we avoid allot of complexity in the model. This allows us to use the model purely to focus on the effect of having a rate-limiting step after equillibrated binding and assembly. Combining **Equations 5-7** yields

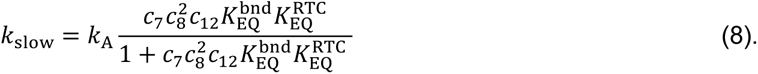

After rearranging the core RTC into an elongation-competent conformation, the protein complex can still potentially unbind from the RNA before incorporating the first NTP (**Figure S6**). The above used that this processes is much faster than the conformational change (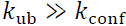). However, this assumption no longer holds true when competing with NTP incorporation (which occurs at *k*_NTP_∼ 70 nt/s) (26). Instead of assuming equilibrium, we solve for the fraction of extended RNA molecules (*P*_NTP_) from the flux-balances of RNA species after the core RTC changed conformation (**Figure S6**):

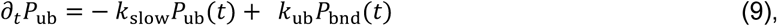

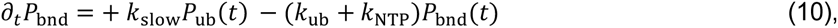

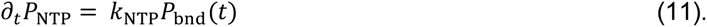

Here, *P*_ub_(*t*) and *P*_bnd_(*t*) represent the time-dependent fractions of RNA molecules (not) bound by a core RTC stabilized in its active conformation. The concentration of RNA molecules on the flow cell’s surface (picomolar range) is negligible when compared to the amount of protein available in solution. A core RTC that unbinds from the RNA is therefore replaced by a different complex that must still undergo the conformational change. Hence, the bare RNA enters the state of being occupied by an activated core RTC at ‘an effective binding rate’ of *k*_slow_ (**Figure S6, Equations 10-11**). Incorporation of the first nucleotide is treated as irreversible (**Figures 5 and S6**). Therefore, any additional number of RNA molecules that are extended between times *t* and d*t* also equals the probability of having started NTP incorporation within that time frame; the first-passage time distribution.

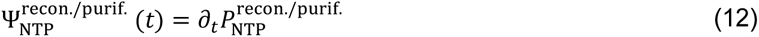

The superscripts ‘recon.’/’purif.’ are used to distuingish between the solutions to **Equations 9-11** under the initial conditions appropriate to the reconstituted/purified core RTC systems respectively. The reconstituted core RTC predominantly bound the RNA before changing conformation (**Figure 5 and Equation 5**, 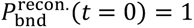, 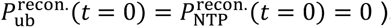. The inverse transition rate into the elongation phase equals the average first-passage time,

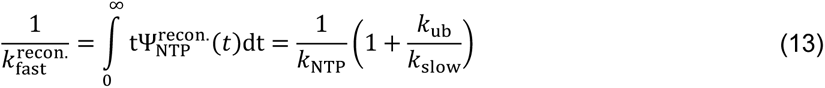

### Modeling the activation time of the purified core RTC

Given a purified complex has already adopted the proper elongation-competent conformation, its activation time only includes the fast component,

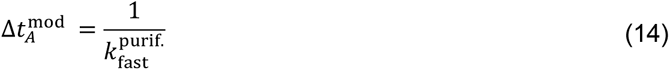

Following the same strategy as done for 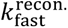, we once again determine the mean first passage time to incorporate the first nucleotide. Different from the case presented above, co-translated complexes in solution already are activated and thus rebind with an intrinsic binding rate (*k*_bnd_). Assigning the same affinity of the purified and reconstituted RTC to the RNA (at a reference concentration of 1 µM), we invoke the law of mass action again,

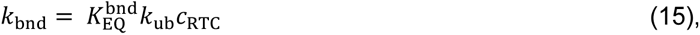

where we used 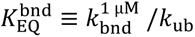; **Figure S6**. Replacing *k*_slow_ by *k*_bnd_ in **Equations 9-10**, results in the inverse reaction rate under the initial condition that the complex must first bind the RNA 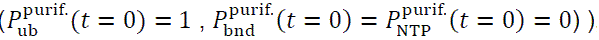,

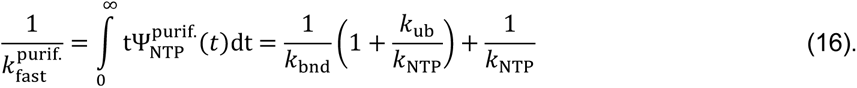

### Incorporating force-dependent binding

At equilibrium, the fraction of bound complex follows the Boltzmann distribution

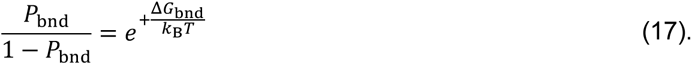

Here, we assigned a free-energy difference for binding the (assembled) RTC on the RNA (Δ*G*_bnd_). The magnetic tweezers apply a constant force (F) to the template strand along the vertical direction (*Z*-direction), thereby putting work into the system. Interactions with the template strand favor binding at lower force (**Figure 2F**), i.e. 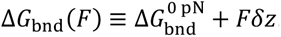, with δ*Z* the length change induced to the RNA by binding of the RTC. Defining a characteristic force of F_0_ = *k*_*B*_*T*⁄δ*Z*, **Equation 7** and **Equation 17** imply a force-dependent equilibrium constant,

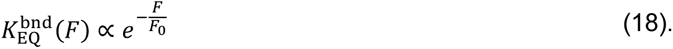

**Equation 18** was applied in **Equation 13** and **Equation 16** to determine force-dependent activation times. The model parameter 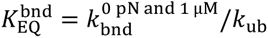 is the constant of proportionality in **Equation 18**, denoting the equilibrium constant in the absence of an applied force. For notational convenience, the equations above still contain 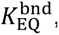, where the force-dependence (**Equation 18**) is implied.

### Fitting procedure using simulated annealing

The Simulated Annealing algorithm (37) is a commonly used algorithm for high-dimensional optimization problems. We used a custom-built Python code that has been more extensively described in (38). We optimized the loss-function (χ^2^) with respect to our model parameters log_10_(*k*_conf_/1 *s*), 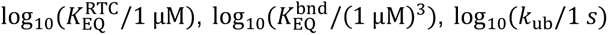, and log_10_(F_0_/1 pN). Model parameters have been transformed to make all fitted parameters lie within the same range.

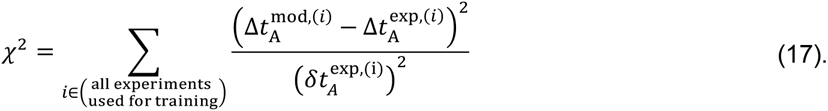

That is, for every primer-extension experiment represented in **Figure 2**, we minimize the sum of differences of our model’s predictions to the corresponding datapoint, weighted by the experimental error 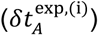 calculated as described above. Trial moves were generated by adding uniform noise of magnitude δ to the present value of each model parameter. The process was initiated with a noise strength of δ = 1.0. In the initiation cycle, the temperature was adjusted until we had an acceptance fraction of 40-60% over 1000 trial moves, based on the Metropolis condition. After this intitial cycle, the temperatures followed an exponential cooling scheme with a 1% cooling rate (*T*_*k*+1_ = 0.99*T*_*k*_). At every temperature, we adjusted the noise strength δ until an acceptance fraction of 40-60% was reached over 1000 trial moves. Once the desired acceptance fraction was reached, an additional 1000 trial moves were performed to allow the system to equilibrate before the next cooling step. Once the temperature had dropped to a factor 10^-4^ of its initial value, we applied the stop condition:

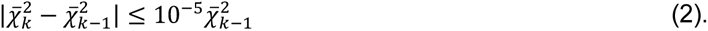

In the above, 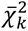 denotes our cost function averaged over the last 1000 trial moves performed at temperature *T*_*k*_. The optimization procedure was repeated 40 times (**Figure S9**). The parameter values with the lowest χ^2^ were used to generate the curves shown in **Figure 2**.

## Results

### Slow activation by reconstituted SARS-CoV-2 core RTCs

To determine the kinetics of the CoV polymerase complex during activation and elongation phases, we built upon our previously developed single-molecule high-throughput magnetic tweezers assay that monitors primer-extension of dozens of individual polymerases simultaneously (26, 34, 35, 39). Magnetic beads were tethered to the glass surface of a flow chamber by a 1043 nt single-stranded (ss) RNA template that included a ∼800 bp primer on the bead proximal side (31) (**Figure 1A** and **Figure S1AB**, **Materials and Methods**). A pair of permanent magnets located above the flow chamber at a fixed height applied a constant attractive force (25 pN, unless specified otherwise) to the magnetic beads that stretched the RNA tethers (32, 40, 41). The three-dimensional position of each bead was tracked in real-time, providing the extension of their respective tether (33). The RNA construct was designed to form a ∼500 bp hairpin when relaxed. The clear signature of the hairpin opening when rapidly increasing the force was used to select for tethers with a properly presented primer (**Figure S1CD**). During the elongation phase, the core RTC converted the ssRNA template into double-stranded (ds) RNA, which decreased the extension of the tether (26).

Initial primer-extension assays were performed using a core RTC reconstituted from nsp7, nsp8 and nsp12-polymerase individually expressed and purified in *Escherichia coli* (*E. coli*) (**Figure 1B, Materials and Methods**). We can define the activation time Δ*t*_A_ as the time from injecting the core RTC components and NTPs until the start of the elongation phase when the tether extension starts decreasing (**Figure 1AC, Materials and Methods)**. These single-molecule experiments showed that the median activation time for the reconstituted core RTC (∼5-10 min) greatly exceeded the elongation time (∼20-40 s) (**Figure 1CD**). To uncover the process(es) giving rise to the observed activation times, we utilized a combination of different strategies to express and/or reconstitute the core RTC.

To perform primer-extension experiments with a core RTC expressed, processed and assembled in the cell, we expressed the complex in *Sf9* cells from a bacmid encoding for nsp12-polymerase, nsp8, nsp7, as well as nsp5-protease, and pulled down and purified the fully assembled complex (hereafter: ‘purified complex’) (**Figure 1B**, **Materials and Methods**). Strikingly, the typical activation time decreased to mere seconds when repeating the experiment using a comparable concentration of purified complex (**Figure 1DE**). We note that all experiments in **Figure 1** were performed at saturating concentrations of core RTC proteins (see below for discussion on **Figure 2**).

Given that individual nsps were expressed in *E. coli* and the purified complex was expressed in *Sf9* cells (**Figure 1B**), we first evaluated whether the different expression system could promote the purified complex’ activation. To this end, we repeated our single-molecule experiments while reconstituting the core RTC with an nsp12-polymerase expressed in *Sf9* cells (**Figure 1BF**, **Materials and Methods**). The resulting reconstituted core RTC, with nsp12-polymerase expressed in insect cells, did not activate rapidly (**Figure 1DF**). We concluded that the difference in expression systems cannot account for most of the observed differences in activation times between bacteria– and insect-derived polymerases.

### Assembly and RNA binding of the core RTC cannot account for the observed activation times

As opposed to the purified complex, the individual proteins of the reconstituted core RTC must come together to assemble into a complex. We questioned whether the time required to assemble the reconstituted core RTC can explain its slower activation as has been suggested in literature (23, 24). To this end, we performed primer-extensions under varying concentrations of one of the three nsps, while keeping the other two unchanged. Here, we used the nsp12-polymerase expressed in *E. coli* to reconstitute the core RTC. While the average activation times measured responded to the varying protein concentrations, we noticed a stronger effect on the fraction of RNA primer-templates that got extended during our 30 min recording (**Figure S2**). This indicated a significant and varying proportion of events (up to ∼75% at the lowest nsp concentrations) were missed as they took longer than the duration of our experiment. The estimated (effective) activation times shown in **Figure 2A-D,F** are maximum likelihood estimates accounting for the fraction of events that exceed the observation time (**Materials and Methods**).

Elongation started sooner on average at higher concentrations of nsp7, with average activation times dropping by ∼4 fold over the tested concentration range (**Figure 2A**). However, at saturating nsp7 concentrations (above 1.8 µM), a substantial effective activation time of ∼1000 s remained (**Figure 2A**), i.e. still significantly slower than the purified complex (**Figure 1E**). A similar trend was seen when increasing either the nsp8 or nsp12-polymerase concentration (**Figure 2BC**), i.e. the activation time decreased until an average time of ∼1000 s remained at saturation, meaning for concentrations of nsp8≥1.8 µM and nsp12-polymerase≥0.2 µM. The remaining time can therefore not be explained by the need to bring the complex together.

To evaluate whether the time required to find and bind the RNA accounts for the remaining activation time, we fixed the stoichiometry (nsp12:nsp7:nsp8 of 1:9:9) and varied the overall concentration both with nsp12-polymerase expressed in bacterial and insect cells (**Figure 2D**). In both cases, an effective activation time of ∼1000 s remained, even above saturating proteins concentration, showing that this time is both independent of host-specific post-translational modifications on the nsp12-polymerase and core RTC assembly, but must be intrinsic to the core RTC components. On the other hand, when we varied the concentration of the purified complex, an activation time of mere seconds was reached at saturation, i.e. activity was detected while flushing in reaction buffer with all proteins into the flow chamber (**Figure 1E** and **Figure 2E**). The saturating concentration of the purified complex is in the tens of nanomolar range (**Figure 2E**), while hundreds of nanomolar of the reconstituted core RTC were required to saturate activity (**Figure 2D**). As the purified complex only needs to find the primer, a lower saturation concentration suggests that the affinity of the complex to the RNA is much greater than that of the proteins to themselves (at the stoichiometry used).

The force applied by the magnetic tweezers provides an additional gauge of the affinity of the core RTC for the primer-template. Namely, elevated tension in the template strand (above ∼35 pN) destabilizes the primer-template junction, hindering a proper placement of the terminal base in the polymerase’s active site upon binding (26). Furthermore, applying tension probes possible conformational changes of core RTC along the direction of the force (e.g. the upstream RNA contacts made by the nsp8 tails (18, 20)), and any conformational change orthogonal to the direction of the force remains unaffected. While we observe a steep increase in effective activation time at forces beyond 40 pN, the effective activation time remains ∼1000 s at lower forces (**Figure 2F**). Taken together, neither the time needed to assemble the core RTC nor the time spent on binding to the RNA can fully account for the observed activation times. We thus conclude that another process must take place to enable the reconstituted core RTC to enter the elongation phase.

### A slow conformational change renders the core RTC elongation-competent

As the duration of the activation phase still greatly exceeded that of the elongation phase at lower forces (**Figure 2F**), we reasoned that the remaining time should be accounted for by processes other than direct interactions along the template RNA. If a slow conformational change is to occur after the core RTC assembled, providing sufficient time for the nsps to interact should induce rapid activation upon introduction of the nucleotides. We pre-incubated nsp12-polymerase with nsp7 and nsp8 for over four hours prior to conducting the single-molecule primer extension assay (**Figure 3A**). To counteract any loss of active proteins during incubation, proteins were added together at 100-fold the concentration used in the flow chamber during the experiment (**Materials and Methods**). Rapid activation was indeed observed with pre-incubated nsp7, nsp8 and nsp12-polymerase individually expressed in *E. coli* (**Figure 3A**). Given sufficient time, reconstituted core RTCs can show the same phenotypical short activation times as the purified complex. However, in experiments performed without any pre-incubation up to ∼80% of core RTCs are capable of activating within 600 s (**Figure S2**). Incubating the nsps for 10 min (nsp12-polymerase expressed in *E. coli*) did not result in any rapidly activating core RTCs (**Figure S3**). The core RTC reconstituted with nsp12-polymerase expressed in *Sf9* did not show rapid activation even after incubation with the co-factors for over 6.5 hours (**Figure 3A**). If the purified complex only activated rapidly because the proteins resided together in the same cell for hours, what then allowed the reconstituted core RTCs to activate within 10 min during our experiments? We surmised that access to the template RNA may have aided the reconstituted system. Introducing the nsps to the RNA in the flow chamber for 10 min indeed resulted in rapid activation upon addition of nucleotides (82% of events using nsp12-polymerase expressed in *E. coli;* 95% of events using nsp12-polymerase expressed in *Sf9*) (**Figure 3B, Materials and Methods**). Taken together, independent of the expression system, pre-incubating the nsps with the RNA reproduced the rapid activation observed with the purified complex (**Figure 3B**). The activation time of ∼1000 s that remained at saturating protein concentrations (**Figure 2A-D**) can thus be accounted for by a process that took place during the incubation with the RNA (**Figure 3AB**). Given the majority of this time is neither spent on assembling the core RTC nor binding the RNA, we concluded that the core RTC must undergo a conformational change that renders it elongation-competent. While the protein complex, at least in part, rearranges itself orthogonally to the RNA template (**Figure 2F**), interactions with the template RNA greatly stabilize the elongation-competent conformation (**Figure 3B**).

### The conformational change enabling elongation requires both nsp7 and nsp8

Having established the existence and necessity of a conformational change for RNA synthesis activity, we next questioned whether nsp7 and nsp8 facilitate it. If the co-factors are a necessary condition, allowing only nsp12-polymerase to access the RNA for 10 min should not result in the rapid activation as seen in the experiments represented in **Figure 3B**. As expected, adding nsp12-polymerase alone with NTP to the RNA did not result in any activity (**Figure 3C**). We subsequently removed all free-floating proteins and thereafter injected the reaction buffer containing nsp7, nsp8 and NTP (**Materials and Methods**). The newly added co-factors can only form a core RTC if a nsp12-polymerase is still bound to an RNA template after rinsing the flow chamber. We successfully recovered activity, but only after approximately 10 min (**Figure 3C**). The nsp12-polymerase preincubated with the RNA template was not yet elongation-competent, but required co-factors to enact the needed conformational change.

To establish whether both nsp7 and nsp8 are needed to induce the conformational change we repeated the experiment described in **Figure 3C**, though this time we added one of the two co-factors together with nsp12-polymerase during pre-incubation with the RNA in the flow chamber (**Figure S4**). Not only are both co-factors required to activate the core RTC, as no activity was recorded after incubation despite the presence of NTP (**Figure S4**), initially omitting one of them severely lowered or even abolished the primer-extension activity. Namely, nsp12-polymerase incubated with nsp7 and the RNA template could only rarely be activated through addition of new nsp7 and nsp8 (∼5% activity vs ∼50% activity when incubating the three proteins with the RNA). Moreover, in the rare cases of activation observed (4 events in total, ∼80 tethers surviving the full duration of the measurement) it was only after an activation time of more than an hour (**Figure S4A**). Allowing nsp12-polymerase and nsp8 to preincubate with the RNA template completely blocked the RTC from activation (**Figure S4B**). No activity was observed within an hour from the point at which all nsps were present (**Figure S4B**). These results suggest nsp12-polymerase explores a conformation space, only to be locked into a stable conformer by the co-factors nsp7 and nsp8. When only one of the two co-factors is available, the polymerase appears to adopt a conformation that prevents RNA synthesis. Escaping this state requires significantly more time (**Figure S4**).

### The elongation-competent conformation of the core RTC is stable for several hours, but is reversed upon loss of nsp7 and nsp8

To establish if a core RTC stably resides in its active conformer, we pre-incubated the purified complex for over eight hours in reaction buffer and at room temperature before performing primer-extensions (**Materials and Methods**). We still observed rapid activation on ∼25% of all tethers upon NTP addition (**Figure S5**), indicating that a substantial fraction of the complexes were still elongation-competent. To test if some of the tethers showed no activity due to disassembly of a bound core RTC, we waited for an additional 10 min, rinsed the flow chamber, and introduced new nsp7, nsp8 and NTP in an attempt to rescue any nsp12-polymerase bound without co-factor (**Figure S5**). Re-introduction of new nsp7 and nsp8 led to new activity (**Figure S5**). Together with the reduction in saturating concentration for the purified complex (i.e. pre-assembled) discussed above, **Figure 2DE**), these results indicate that nsp12-polymerase binds the RNA template with a higher affinity than nsp7 and nsp8 bind to nsp12-polymerase. A long-lasting activation phase was observed again, indicating that the core RTC reverts to its inactive form upon loosing co-factors (**Figure S5**).

Taken together, the core RTC must undergo a conformational change to initiate RNA synthesis (**Figure 3AB**). This conformational change requires both nsp7 and nsp8 to be present during the assembly process to prevent the complex from being stabilized in an inactive form (**Figure 3C** and **Figure S4**). After successfully having undergone the correct transition, the core RTC is stably locked into an active conformation for several hours (**Figure S5**).

### Dynamics during the elongation phase are independent of those during the activation phase

Next, we set out to determine whether the long lag-times prior to detectible primer-extension in bulk (23, 24) are completely explained by the time required to escape the activation phase. To this end, we examined the dynamics during the ensuing elongation phase. Our single-molecule experiments allow us to directly tell if an increased yield of product RNA is due to more core RTCs successfully entering the elongation phase, or due to an enhanced rate of RNA synthesis. During the elongation phase, the tethered magnetic bead moved downwards (**Figure 1AC**). Converting the height drop to the fraction of ssRNA that has been converted into dsRNA gives rise to the position of the core RTC along the template (34) (**Figure 4A**). Comparing the purified complex and reconstituted core RTCs, we observed no discernible difference amongst their time trajectories (**Figure 4A**). Given the rich, stochastic, variation amongst traces within an experiment we quantified the duration of the elongation phase (Δ*t*_E_) together with the resulting length of the product (Δ*L*_E_) (**Figure 4AB**). No significant difference in either quantity was detected amongst the three expression systems tested (**Figure 4B**). To completely characterize the dynamics of the obtained time traces we further extracted the distribution of times required to add ten consecutive nucleotides (11, 26, 35, 39, 42). None of the three dwell time distributions shown in **Figure 4C** were significantly different from the distribution we previously reported to hold true for a (reconstituted) core RTC (26). The pause-free elongation rate can be estimated by the location of the main peak seen in the histograms (26), and the three of them coincide in **Figure 4C**. We found no evidence for any influence of the choice of expression system on the core RTC’s dynamics during the elongation phase. As the activation times increased with decreasing co-factor concentration(s) (**Figure 2AB**), we examined the elongation phases under varying nsp7 (**Figure 4D-F**) or nsp8 (**Figure 4G-I**) concentrations. The concentration of co-factors in solution had no influence on elongation times, product lengths, or the dwell time distributions. All measured distributions coincide with our previously reported one (11, 26).

We conclude that any lag-time before RNA production seen in bulk biochemistry experiments (23, 24) stems entirely from altered dynamics during the activation phase. Our previous model of the elongation phase in the reconstituted sytem (26) should therefore hold true irrespective of the dynamics of the activation phase.

### Activation of the core RTC follows its equilibrated assembly

To further test if the existence of a conformational change after core RTC assembly can quantitatively describe our data we built a kinetic model based on our above findings. **Figure 5** shows the predominant assembly and activation pathway of the core RTC towards processive RNA synthesis. The complete reaction pathway underlying the kinetic model fitted to the data is shown in **Figure S6 (Materials and Methods)**. The core RTC is able to sequentially assemble on the RNA, starting with nsp12-polymerase, followed by the addition of nsp7 and nsp8 (**Figure 3C** and **Figure S5**). Both co-factors (nsp7 and nsp8) must be present to obtain active core RTCs to any appreciable degree (**Figure S4**). Furthermore, the core RTC activates within minutes in the presence of RNA, whereas the apo form, i.e. without RNA, requires several hours of incubating the nsps to see rapid activation (**Figure 3AB** and **Figure 5**).

We observed long activation times of several minutes, even at saturating protein concentrations (**Figure 2A-D**), indicating that activation occurs after assembly of the core RTC onto the RNA, which in itself can be treated as equillibrated within that timespan (**Figure 5** and **Figure S6**). For each experiment, the distribution of activation times was consistent with a single exponential with a characteristic timescale dependent on protein concentrations (**Figure S7**) and force (**Figure S8**). Hence, we conclude that a single rate-limiting conformational change is responsible for the activation of the core RTC (indicated with the rate *k*_conf_ in **Figure 5** and **Figure S6**). The purified complex showed rapid elongation upon addition into the flow chamber (**Figure 1E**), suggesting this complex had already gone through activation, likely during protein expression. Furthermore, the activated conformation of the core RTC proved to be reasonably stable for over 8 hours of pre-incubation (**Figure S5**). We therefore modelled the activation step as irreversible (**Figure 5**, **Figure S6**). Having adopted the elongation-competent conformation, the activation time is only limited by the rate of nucleotide incorporation (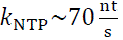 at 25 °C, **Figure 4**) (26). It took micromolars of (co-)factors (**Figure 2A-C**, **Figure S2C-F**), while just tens of nanomolar of the purified complex was needed to saturate activity (**Figure 2E** and **Figure S2I-J**). Furthermore, activity can be restored on RNA initially bound by nsp12-polymerase alone by adding new nsp7 and nsp8 (**Figure 3C** and **Figure S5**). Hence, we assume that the core RTC binds much tighter to the RNA than the complex is held together (**Figure S6**, **Materials and Methods**). The kinetic model that we fitted to the data globally (**Figure S6**, **Figure 2**, **Materials and Methods**) quantitatively captures all experiments in a unified manner (**Figure S9** shows the distribution of fit parameters with fit results within 1% of the best fit, **Materials and Methods**). Although the model allows for multiple cycles of core RTCs binding and activating prior to starting RNA synthesis (**Figure S6**, **Materials and Methods**), we find a good agreement with our data when the unbinding rate (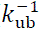 between 1 – 20 s, **Figure S9**) is much lower than the rate of nucleotide addition (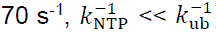). Taken together, following equilibrated assembly (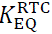 between 0.06 – 0.99 μM^−4^, **Figure S9**) and RNA binding (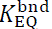 between 60 – 980 μM^−1^ at 25 pN, with a characteristic force F_0_ between 7.6 – 8.2 pN, **Figure S9**), the rate of starting RNA synthesis is trully limited by the time required for the first core RTC to undergo the conformational change (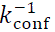 between 773 – 800 s, **Figure S9**).

### SARS-CoV-2 core RTC activation remains slow in the presence of nsp13-helicase

During a viral infection, the core RTC associates with additional nsps to form an extended RTC, such as nsp13-helicase (15, 43). We addressed whether the associated with nsp13-helicase to the core RTC results in a similarly large barrier towards elongation-competence. We performed primer-extension experiments with the three core RTC proteins at saturating concentrations (0.2 µM nsp12-polymerase, 1.8 µM nsp7, and 1.8 µM nsp8), as well as purified nsp13-helicase at the indicated concentration (**Figure S10AB, and Materials and Methods**). Structural studies have reported that two nsp13-helicases bind the core RTC, one of them also binding the template RNA (15). Accordingly, we see a reduction in activation times upon addition of nsp13-helicase (**Figure S10C-E**). While an increase in tension lowered the affinity of the core RTC to the RNA (**Figure 2F**), addition of nsp13-helicase increased its affinity. We further note that in the presence of saturating amounts of nsp13-helicase, i.e. above 10 nM, the duration of the activation phase is still set by one characteristic timescale (**Figure S10C**). Moreover, the addition of nsp13-helicases only minimally increased the success rate of the primer-extension reaction (**Figure S10F**). In terms of our kinetic model, the helicase increases both the RNA-binding affinity (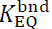) and transition rate into elongation-competency (*k*_conf_). Regardless, an average activation time of ∼360 s remains at a saturating concentration of 20 nM nsp13-helicase (**Figure S10C-E**). This demonstrates that the time needed for a conformational change identified for the core RTC remains relevant when viewed in the context of the extended complex.

## Discussion

The coronavirus RTC, in particular the ‘core RTC’, fulfills the essential role of replicating the viral genome (13) and is therefore a key target for antiviral drugs. There is a wealth of structural (8, 10, 18, 20) and biochemical studies focused on understanding and inhibiting the CoV polymerase’s RNA synthesis capabilities (9, 11, 12, 26, 28, 44). The assembly of the core RTC was earlier proposed to be the rate-limiting step in the RNA synthesis reaction (23, 24). The present study reveals that an already assembled core RTC must undergo a conformational change that represents the true rate-limiting step towards processive elongation.

We expanded the single-molecule high-throughput magnetic tweezers assay we previously developped (26) to assess the lag time between injection of the RTC proteins into the flow chamber and the start of RNA synthesis (**Figure 1A**). We investigated how varying the stoichiometry and concentration of the core RTC proteins, as well as their mode of expression, impacted both activation and elongation phases. We showed that the approximately ten minutes activation time (**Figure 1**) did not result from either the time required for the core RTC to assemble or to bind the RNA (**Figure 2**), but rather from a conformational change within the complex (**Figures 3**, **Figure 5**).

We note that when expressing nsp12-polymerase in *E. coli*, the gene sequence was codon optimized not for total protein production but to maximize activity in bulk (27, 45) (**Materials and Methods**). Rare codons were intentionally maintained within the coding sequence with the idea of retaining natural pause sites for the ribosome to allow the protein more time to fold into a functional conformer during translation. While clearly not an elongation-competent conformer from the onset (**Figure 1C**), an altered free-energy landscape for folding the nsp12-polymerase can facilitate access to the elongation-competent conformer. This likely explains why pre-incubating the *E. coli* expressed nsp12-polymerase with nsp7 and nsp8 resulted in rapid activation, while repeating the same experiment with the *Sf9* expressed nsp12-polymerase whose coding sequence was codon-optimized for protein yield, i.e. lacking the rare codons, did not (**Figure 3A**). It also supports our hypothesis that nsp7 and nsp8 stabilize a conformation in which nsp12-polymerase properly exposes its active site to the RNA template.

Activation of the SARS-CoV-2 core RTC requires the co-factors nsp7 and nsp8 (**Figure 3**, **Figure 5**). Cryogenic-electron microscopy (cryo-EM) studies of apo-core RTC and RNA bound core RTC revealed nsp8 must undergo a large conformational change allowing its N-terminal tails to form rigid contacts ∼28 bp upstream of the active site (18–20). While this is clearly a conformational change required for RNA synthesis, our data does not support the interpretation that the rotation of the nsp8s constitutes the activation step. Namely, using a purified core RTC or pre-incubating the individual nsps allowed for rapid activation (**Figure 1D**, **Figure 3B**). In these core RTCs, the nsp8 monomers must still unfold their N-terminal tails prior to starting RNA synthesis (17, 19). Given the nsp8 tails align along the RNA, establishing the contacts should be tension-dependent. Furthermore, we saw a reduction in the mean activation time when including nsp13-helicase (**Figure S10**), suggesting that nsp13-helicase successfully bound the extended nsp8-tails (15). Finally, varying the tension applied to the template RNA did not reduce the (effective) activation time to seconds (**Figure 2F**). Therefore, the activation step we reveal here is a distinct conformational change that, at least in part, rearranges protein domains that either do not interact with the RNA or move orthogonal to the force. We hypothesize that the conformational change is mainly in nsp12-polymerase, with the co-factors stabilizing a conformer in which the RNA is positioned properly for RNA synthesis. In the absence of either nsp7 or nsp8, nsp12-polymerase is forced into a conformation that is inactive and is harder to escape from (**Figure S4**).

Previous studies have identified viral genomic RNA-polymerase interactions to initiate either replication or transcription in RNA viruses. The dengue virus polymerase (NS5) interacts with the stem loop 5 in the viral genome, which enables its transition into an ‘elongation complex’ (46). Alphaviruses have a promotor sequence in their genome that activates the RdRp to transcribe a subgenomic RNA encoding for structural proteins (47). Our results also indicate an important role for the RNA in activating the core RTC, which was orders of magnitudes faster with RNA as opposed to without (**Figure 3**). The rich structure of the SARS-CoV-2 genome (48) may contain a motif yet to be discovered to recruite and activate the RTC.

Recent studies demonstrated that a P323L mutation in the nsp12-polymerase, present in most SARS-CoV-2 variants of concern (49), increases the core RTC’s primer-extension rate (50) and the viruses’ fitness (51). The mutated residue stabilizes the nsp12/nsp8 interface (50). Based on our results, we suspect the mutation modulates the RTC’s activation time. One may speculate how such a mutation could be beneficial to the virus. Seminal work on CoV murine hepatitis virus showed that stopping translation in infected cells rapidly resulted in a drop of (-)RNA synthesis (52), indicating that a continuous production of nsps was necessary for viral RNA synthesis during infection. Hence, the virus is seemingly assembling and activating new RTCs throughout the infectious cycle. During a viral infection, the extended RTC incorporates additional components beyond the core. These include nsp13-helicases (15), an nsp10-nsp14 complex with exonuclease and methyltransferase activities (16, 53), which may also associate with nsp16 (54), and the capping co-factor nsp9 (27), among others (16). The experiments of **Figure S10** show that the polymerase-helicase complex must still undergo a post-assembly conformational change to become elongation-competent. The extended activation time of the core RTC may enable the recruitment of all essential nsps before replication begins. The association of additional components could influence the kinetics of this activation, a possibility that warrants future investigations.

Our results provide a mechanistic insight into the assembly and activation of the core RTC, establishing a platform to recruit other nsps and assemble a complete and functional CoV RTC to reveal the molecular determinants of CoV replication.

## Acknowledgements

We thank Gonzalo Cosa for fruitfull discussions. DD was supported by the Interdisciplinary Center for Clinical Research (IZKF) at the University Hospital of the University of Erlangen-Nuremberg, the German Research Foundation grant DFG-DU-1872/4-1, BaSyC – Building a Synthetic Cell” Gravitation grant (024.003.019) of the Netherlands Ministry of Education, Culture and Science (OCW) and the Netherlands Organisation for Scientific Research (NWO), NIH fundings R01 AI161841-01, U19 AI171292 and U19 AI171421, and NWO funding OCENW.XL21.XL21.115. IA was supported by NIH funding R01 GM067153. KDR was supported by NIGMS funding R35-GM12260.

## Author contributions

DD, SCB, AD, MK, CEC, JJA designed the research. AD, SCB and MK performed the experiments, as well as processed and analyzed the data. MK and MD developed the kinetic model. BW and IA provided recombinant nsp12-polymerase expressed in *E. coli*. TKA and RNK provided recombinant nsp12-polymerase expressed in *Sf9*, as well as nsp7 and nsp8. DK, HWL and MG provided co-translated core RTC. JCM and KDR provided recombinant nsp13-helicase. DD supervised the research. MK and DD wrote the article. All the authors edited the article.

## Competing interests

The authors declare that they have no competing interests.

## Data availability

*Data will be made publically available upon publication (DOI link will be added)*.

## Notes

### Competing Interest Statement

The authors have declared no competing interest.

