## Supplementary Information for "A post-assembly conformational change makes the SARS-CoV-2 polymerase elongation-competent"

This document contains 10 supplementary figures.

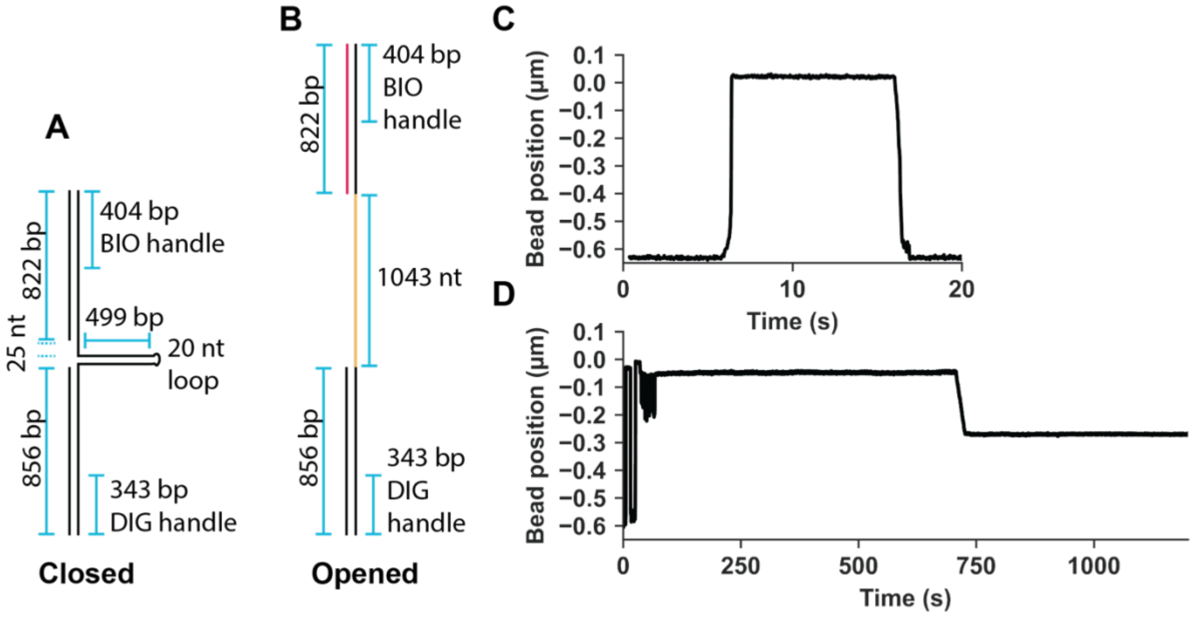


**Figure S1, relates to Figure 1: Test of template validity.** (**A**) Schematic of RNA construct used in magnetic tweezers assay. The construct contains a 499 bp hairpin, and is flanked by biotin- (BIO) and digoxigenin-handles (DIG) for attachment to magnetic beads and the glass surface respectively. A 25 nt gap is included between the BIO-handle and the start of the hairpin (**Materials and Methods**). (**B**) Schematic of the RNA construct when the hairpin is in its open configuration. (**C**) Prior to the primer-extension experiment a short force-extension experiment is run to select for magnetic beads tethered by proper hairpin RNA constructs (Materials and Methods). About 8 seconds into the experiment, we quickly ramp the force up to approximately 40 pN. The presence of a hairpin on the RNA construct is readily detected as a clear ~0.6 µm jump in end-to-end distance once a critical force around 22 pN is reached. After maintaining the highest force for 15 seconds, the force is lowered again, resulting in the hairpin closing and the end-to-end distance dropping again. **(D)** Example primer-extension trace. The same trace is shown (from the addition of the proteins onwards) in **Figure 1C**. Prior to flushing the proteins and NTP into the flow cell, a short force-extension protocol is run to once again to check if the hairpin opened properly.

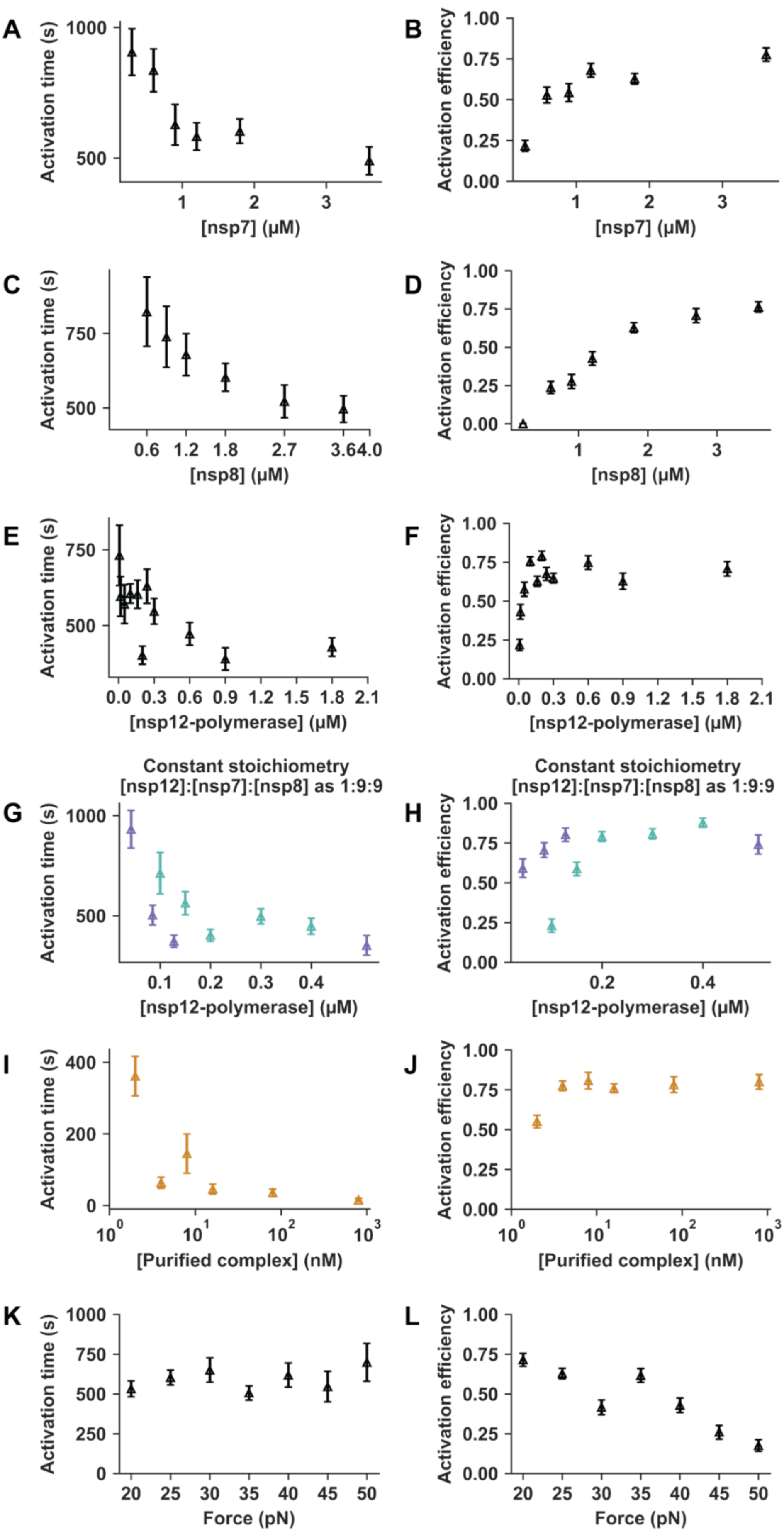
A
B
C
D
E
F
G
H
I
J
K
L
Activation time (s)
Activation efficiency
[nsp7] (μM)
[nsp8] (μM)
[nsp12-polymerase] (μM)
Constant stoichiometry
[nsp12]:[nsp7]:[nsp8] as 1:9:9
[Purified complex] (nM)
Force (pN)

**Figure S2, relates to Figure 2: Recorded activation times and efficiencies.** When not being varied, experiments were performed using 0.16 μM of nsp12-polymerase, 1.8 μM nsp7, 1.8 μM nsp8, and 25 pN. **(A)** Recorded activation times and (**B**) fraction of tethers with properly opened hairpin (**Figure S1**) that display the signature of primer-extension ('Activation efficiency') across nsp7 concentration. Equivalent figures are shown across (**C-D**) nsp8 concentration, (**E-F**) nsp12-polynmerase concentration, **(G-H**) nsp12-polymerase concentration while maintaining a nsp7 and nsp8 concentration that was 9 times greater, (**I-J**) concentration of purified complex, and (**K-L**) force. In A-F, K, and L the core RTC was reconstituted using the nsp12-polymerase expressed in *E. coli*. In G and H, the core RTC was reconstituted using the nsp12-polymerase expressed in *E. coli* (turquoise) or *Sf9* (purple). Error bars represent 95% confidence intervals determined as described in Materials and Methods.

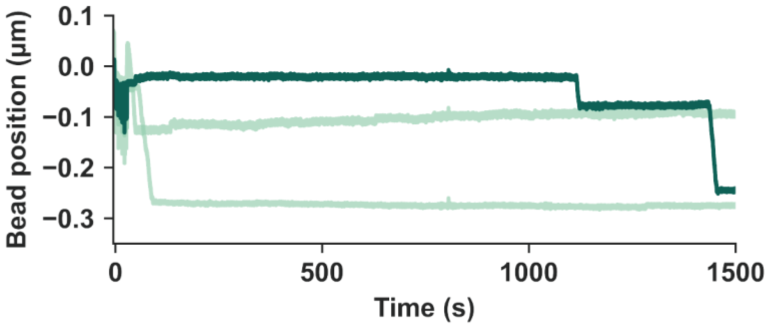


**Figure S3, relates to Figure 3: Ten minutes of incubating reconstituted core RTC.** Example time traces of a core RTC with its nsp12-expressed in *E. coli*. Prior to the measurement proteins have been incubated at 25 ˚C for 10 minutes (following the protocol shown in **Figure 3A**).

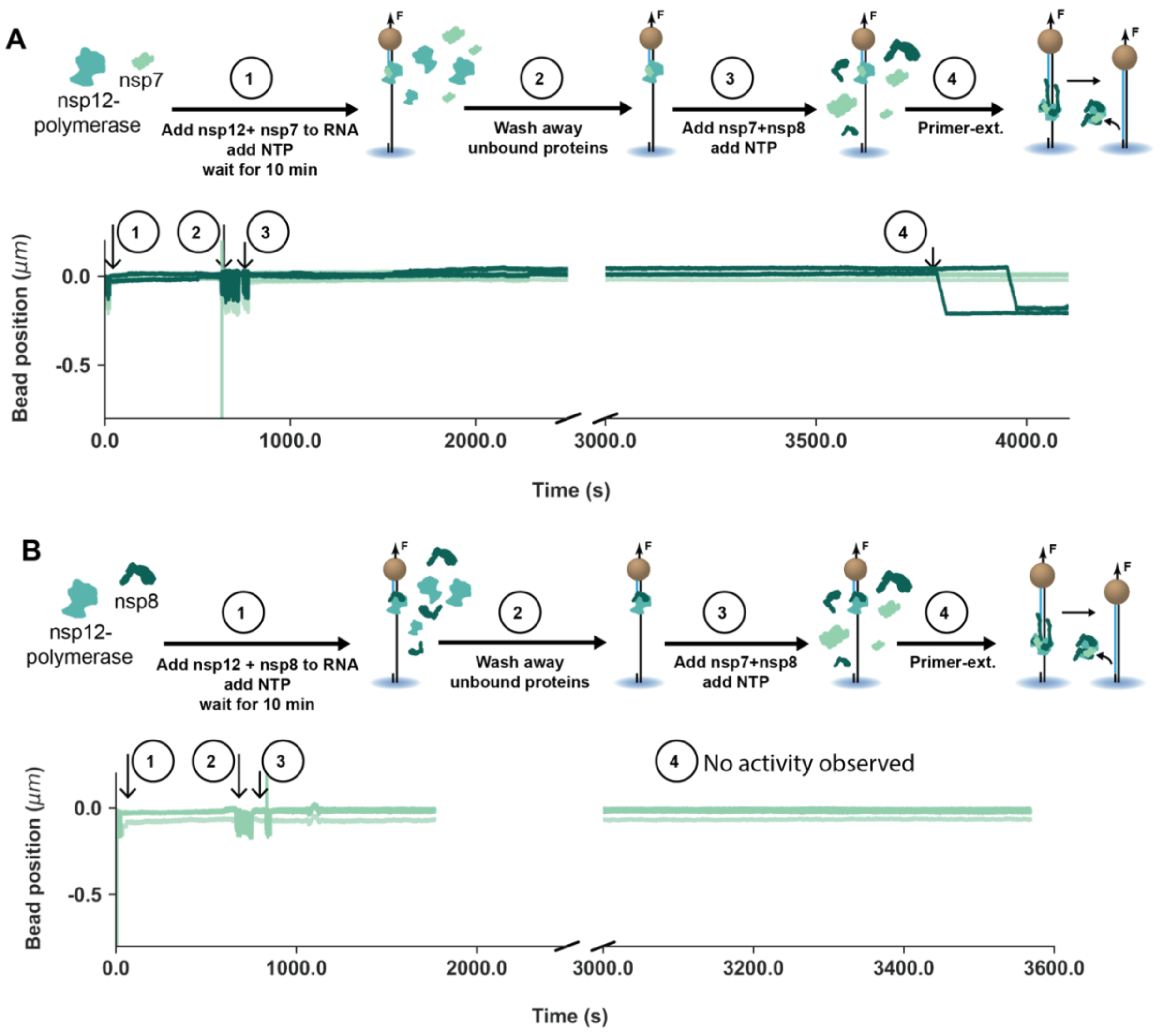


**Figure S4, relates to Figure 3: Both co-factors are needed to activate the core RTC.** Example time traces of a core RTC with its nsp12-expressed in *E. coli.* Prior to measurement, nsp12-polymerase, either **(A)** nsp7 or **(B)** nsp8, and NTP where injected into the flow chamber with the immobilized RNA templates. After ten minutes, the flow cell was cleared from free-floating proteins and nsp7, nsp8 and NTP were added. Rare cases of RNA extension activity were observed using nsp12+nsp7 (A). Using nsp12+nsp8, no activity was observed (B).

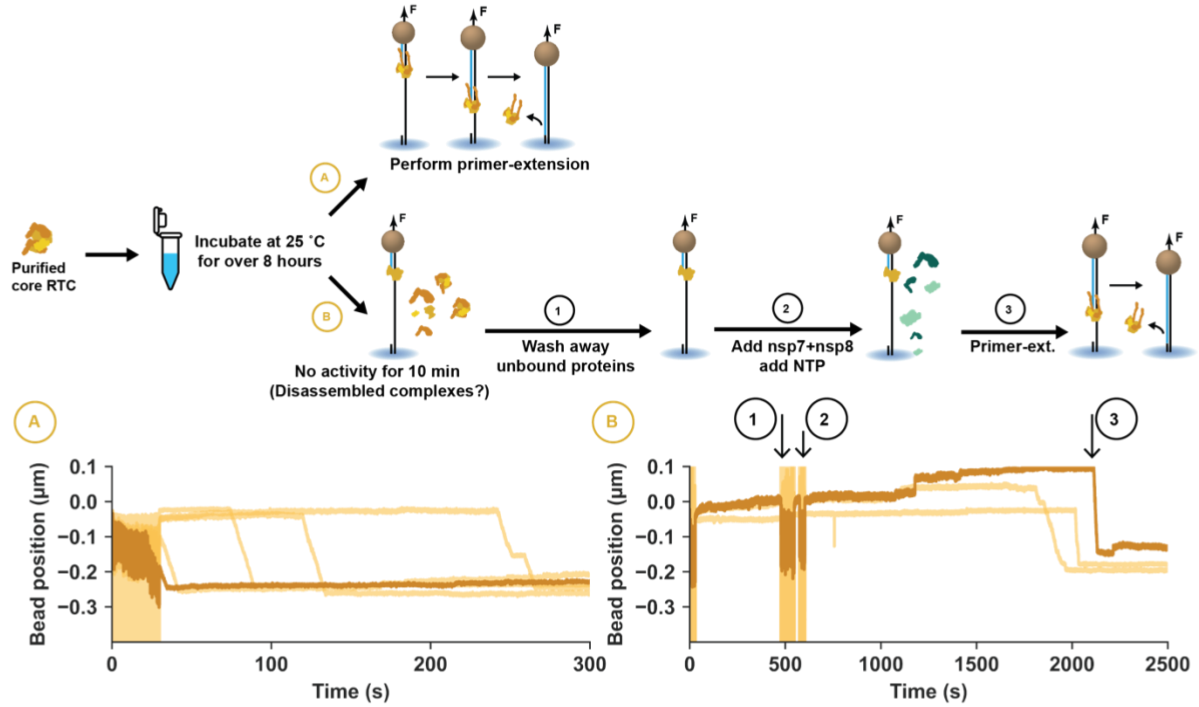


**Figure S5, relates to Figure 3: Pre-incubation of co-translated core RTC.** 80 nM of purified complex was incubated at 25 ˚C for over 8 hrs prior to starting the primer-extension experiments. Of all traces with a properly opened hairpin (see **Figure S1**), 25% displayed rapid onset of the elongation phase as shown in the example traces (path A). Continuing the same recording (path B), excess proteins were removed after 10 minutes (step B.1), and 1.8 µM of nsp7 and nsp8 as well as 500 µM of all four NTP were introduces (step B.2). Example time traces in which activity was recovered (step B.3) are shown.

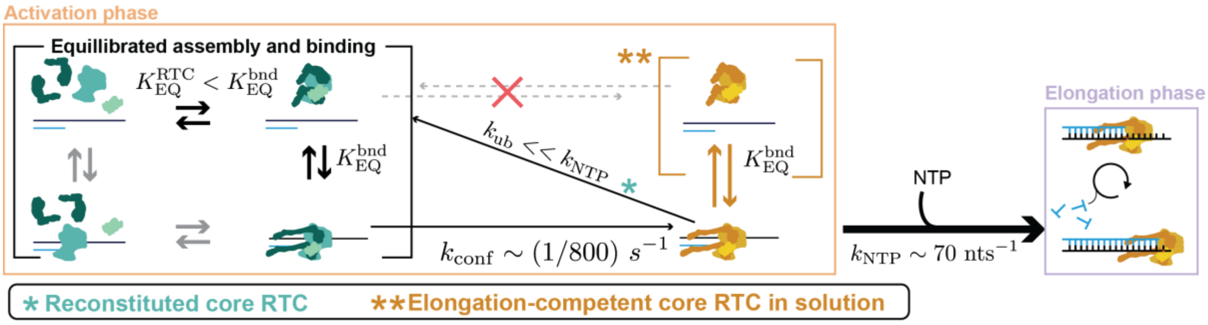


**Figure S6, relates to Figure 5: Kinetic model of core RTC assembly and activation.** Schematic of the kinetic model used fitted against the data shown in **Figure 2**. See **Materials and Methods** for details. Equilibrated assembly of the core RTC from the nsps (equilibrium constant $K_{\mathrm{EQ}}^{\mathrm{RTC}}$) and to the RNA (force-dependent equilibrium constant $K_{\mathrm{EQ}}^{\mathrm{bnd}}(F) \propto e^{-F/F_0}$) is followed by a rate-limiting conformational change ($k_{\mathrm{conf}}$) to make the complex elongation-competent. Undergoing the conformational change in solution is orders of magnitude slower compared to the RNA-bound complex (red cross over right-pointing grey arrow). Similarly, the conformational change is treated as irreversible (red cross over left-pointing grey arrow). Replacing an unbound core RTC requires a new complex to undergo the conformational change when reconstituting the core RTC (turquoise). The purified complex is treated as a pre-activated core RTC. When present in solution, these can directly bind the RNA in their elongation-competent form, which by-passes the activation step (orange).

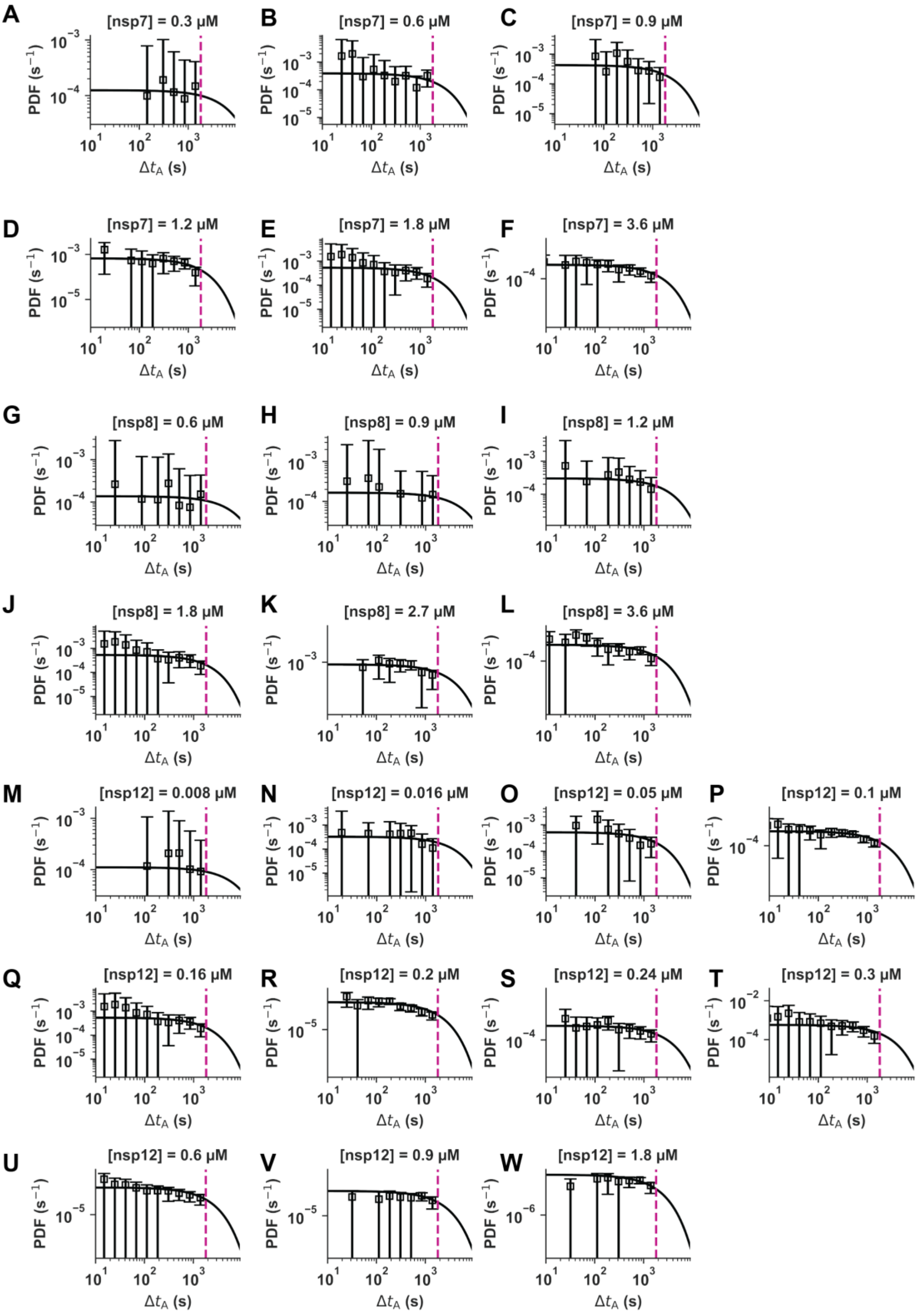


**Figure S7, relates to Figures 2, 5 and S2: A single time scale sets the activation time.** When not being varied, experiments were performed using 0.16 μM of nsp12-polymerase, 1.8 μM nsp7, 1.8 μM nsp8, and 25 pN. Normalized histogram of activation times for the primer-extension experiments under varying **(A-F)** [nsp7], **(G-L)** [nsp8], and **(M-W)** [nsp12]. Error bars represent 95% confidence intervals determined using 1000 bootstrap samples. Maximum likelihood estimation was used to determine the time constants of the single-exponential distribution that best describes the data (**Materials and Methods**). The corresponding distributions (shown as solid lines) have a portion under the curve beyond the duration of our experiment (1800 s, pink dashed line). The corresponding characteristic times are shown in **Figure 2**.

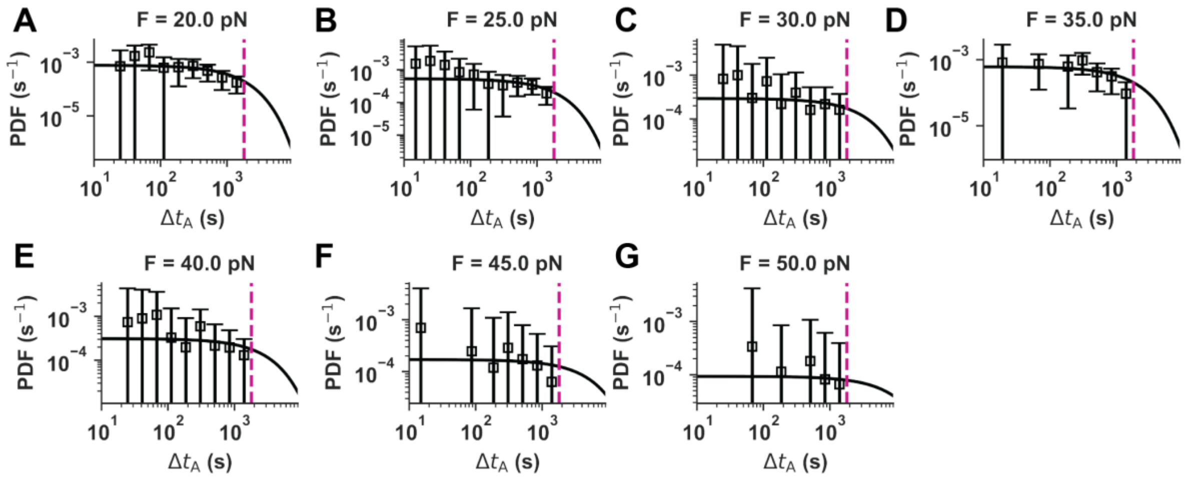


**Figure S8, relates to Figures 2, 5 and S2: A single time scale sets the activation time.** Normalized histogram of activation times for the primer-extension experiments under varying force. Error bars represent 95% confidence intervals determined using 1000 bootstrap samples. Maximum likelihood estimation was used to determine the time constants of the single-exponential distribution that best describes the data (**Materials and Methods**). The corresponding distributions (shown as solid lines) have a portion under the curve beyond the duration of our experiment (1800 s, pink dashed line). The corresponding characteristic times are shown in **Figure 2F**.

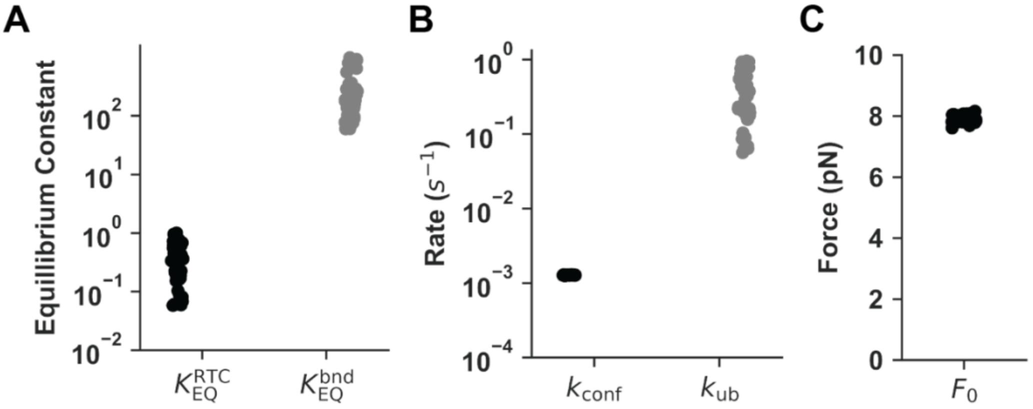


**Figure S9, relates to Figures 5 and S6: Distribution of fit parameters.** Fitted parameter values across the 40 fits performed, all with a final value of the loss function ($\chi^2$, **Materials and Methods**) within 1% of the best fit. **(A)** Equillibrium constants ($K_{\mathrm{EQ}}^{\mathrm{RTC}}$ measured in μM$^{-4}$, $K_{\mathrm{EQ}}^{\mathrm{bnd}}$ at 0 pN of force measured in μM$^{-1}$) . **(B)** rate parameters in 1/s. **(C)** Characteristic force ($F_0$) that determines the sensitivity of $K_{\mathrm{EQ}}^{\mathrm{bnd}}$ to force ($K_{\mathrm{EQ}}^{\mathrm{bnd}} \propto e^{-F/F_0}$ (**Materials and Methods**).

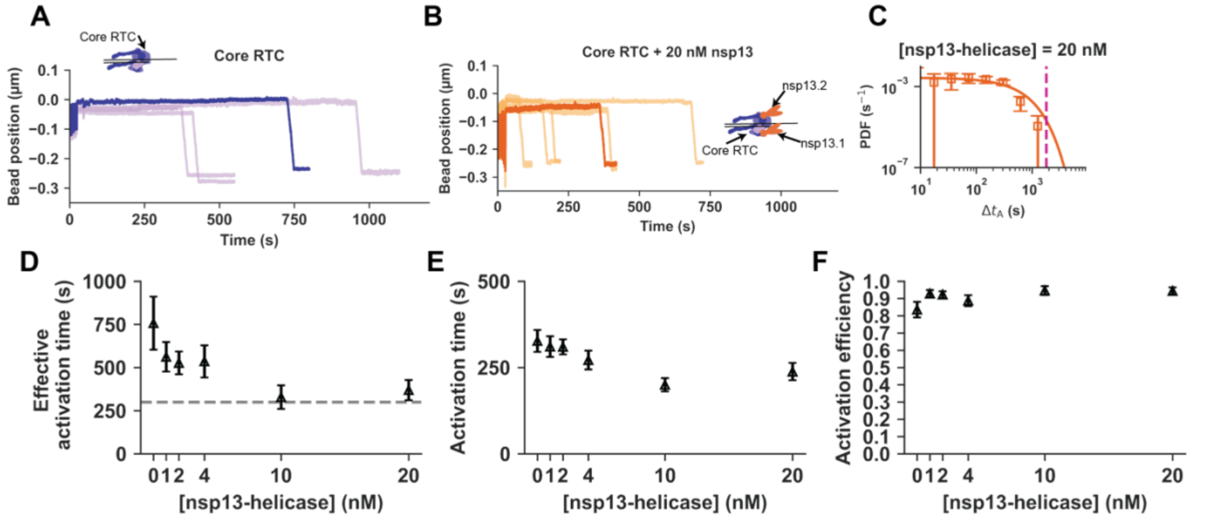


**Figure S10, relates to Figures 2 and 5: Kinetics during the activation phase of the core RTC + nsp13-helicase.** Primer-extensions are performed using the same assay as displayed in **Figure 1A** with 0.16 µM nsp12-polymerase (expressed in *Sf9*), 1.8 µM nps7, 1.8 µM nsp8, and 25 pN. **(A)** Example time traces for the reconstituted SARS-CoV-2 core RTC without nsp13. **(B)** Example time traces for the reconstituted SARS-CoV-2 core RTC + nsp13-helicase. **(C)** Normalized histogram of activation times ($\Delta t_A$) when including 20 nM of nsp13-helicase. Error bars represent 95% confidence intervals determined using 1000 bootstrap samples. Maximum likelihood estimation was used to determine the time constant of the single-exponential distribution that best describes the data (**Materials and Methods**). The distribution (shown as a solid line) has a portion under the curve beyond the duration of our experiment (1800 s, pink dashed line). **(D)** Effective mean activation time (dashed horizontal line is shown to guide-the-eye), **(E)** measured mean activation times, and **(F)** activation efficiency vs nsp13-helicase concentration. In **(D, E, F)**, the error bars represent 95% confidence intervals estimated through bootstrapping (**Materials and Methods**).
